# Fight not flight: parasites drive the bacterial evolution of resistance, not escape

**DOI:** 10.1101/2023.04.29.538831

**Authors:** Michael Blazanin, Jeremy Moore, Sydney Olsen, Michael Travisano

**Affiliations:** University of Minnesota; Yale University; Yale University; University of Minnesota

## Abstract

In the face of ubiquitous threats from parasites, hosts can evolve strategies to resist infection or to altogether avoid parasitism, for instance by avoiding behavior that could expose them to parasites or by dispersing away from local parasite threats. At the microbial scale, bacteria frequently encounter viral parasites, bacteriophages. While bacteria are known to utilize a number of strategies to resist infection by phages, and can have the capacity to avoid moving towards phage-infected cells, it is unknown whether bacteria can evolve dispersal to escape from phages. In order to answer this question, we combined experimental evolution and mathematical modeling. Experimental evolution of the bacterium *Pseudomonas fluorescens* in environments with differing spatial distributions of the phage Phi2 revealed that the host bacteria evolved resistance depending on parasite distribution, but did not evolve dispersal to escape parasite infection. Simulations using parameterized mathematical models of bacterial growth and swimming motility showed that this is a general finding: while increased dispersal is adaptive in the absence of parasites, in the presence of parasites that fitness benefit disappears and resistance becomes adaptive, regardless of the spatial distribution of parasites. Together, these experiments suggest that parasites should rarely, if ever, drive the evolution of bacterial escape via dispersal.

## Introduction

In nature, organisms have numerous evolutionary responses to the ubiquitous threats they face from parasites. One common strategy is resistance, where defenses prevent the establishment and proliferation of parasites (Hall et al. 2017; Gibson and Amoroso 2022). However, another strategy is avoidance, which precludes infection by reducing the rate of contact between hosts and parasites in the first place (Gibson and Amoroso 2022). In fact, examples of parasite avoidance are common. For instance, many organisms will avoid behavior that could expose them to parasites (Stephenson et al. 2018; Paciência et al. 2019), while others will use dispersal as a mechanism to escape the local threat of parasitism (Sloggett and Weisser 2002; Fill et al. 2012; Baines et al. 2020; Brophy and Luong 2021; Zilio et al. 2021).

Spatial factors, like the distribution of parasites in space, are predicted to alter the selection for resistance and avoidance. For instance, selection for resistance strengthens with spatial homogeneity and parasite spread (Brockhurst et al. 2003; Vogwill et al. 2008), while selection for avoidance strengthens as the local risk of infection or spatial heterogeneity of the environment increase (Boulinier et al. 2001; Weisser 2001; Baines et al. 2020).

One domain where host-parasite interactions are extremely common is that of microbes. In particular, bacteria and their viral parasites, bacteriophages, are the two most abundant biological entities on the planet (Whitman et al. 1998; Hendrix et al. 1999; Bar-On et al. 2018; Mushegian 2020). When a phage infects a bacterial cell, it co-ops the cellular machinery to replicate, often harming or killing the host. This host-parasite interaction between bacteria and phages has major effects, from human health (Kortright et al. 2019) to the global ecosystem (Finlay et al. 1997; Whitman et al. 1998; Wommack and Colwell 2000; Weitz and Wilhelm 2012).

In the face of constant threats from phages, bacteria have been found to utilize a number of strategies to resist infection by phages (Hampton et al. 2020). For instance, bacterial populations can evolve resistance by modifying the cellular structures that phages rely on, especially the extracellular structures used by phages to initially attach to the cell. Bacteria can also carry any number of cytoplasmic defenses that operate after a phage infects the cell. In fact, bacteria can even activate intracellular defenses in response to signals from phage-infected neighbors (Høyland-Kroghsbo et al. 2013, 2017; Tan et al. 2015; Patterson et al. 2016; Baskerville et al. 2018; de Mattos et al. 2022). Moreover, recent advancements have revealed an enormous diversity of previously-undiscovered defense systems (Millman et al. 2020, 2022), suggesting that many more bacterial responses to phages remain to be found.

While bacterial resistance to phages has been extensively studied, bacterial avoidance of phages has not. Many bacteria do have complex motility behaviors that they use to disperse and physically navigate their environment (Wadhwa and Berg 2022). For instance, bacteria can detect a number of attractant and repellant signals. They also have several mechanisms for motility, including tail-like flagella that enable swimming or swarming in liquid and semi-solid environments, pili for twitching motility on surfaces, and extracellular adhesions for gliding across surfaces (Wadhwa and Berg 2022). However, while the molecular mechanisms of many of these systems are well-established, the ecological functions they fulfill in natural environments, including whether they enable avoidance of phages, often remain unresolved (Keegstra et al. 2022).

To our knowledge, there is only a single example of bacterial avoidance of phage infection (Keegstra et al. 2022). Bru et al. found that bacteria will avoid swarming towards molecules signaling stress, including those expressed during phage infection (Bru et al. 2019). This shows that bacteria can avoid behavior that could expose them to parasites. However, it remains unclear whether bacteria could evolve dispersal to escape away from a local threat of parasitism. Some compelling work has indirectly tested this idea in spatially structured environments. For instance, Li et al. and Taylor & Buckling carried out studies on dispersing bacteria and phages. Their ecological data supported the evolutionary prediction that phages could select for bacteria with greater dispersal (Taylor and Buckling 2013; Li et al. 2020). In contrast, the ecological and evolutionary experiments of Ping et al. implied that the presence of phages should select for resistance and not dispersal (Ping et al. 2020).

Here, we carry out the first direct test of the hypothesis that bacteria can evolve dispersal to escape phages. To test this hypothesis, we manipulated the spatial distribution of phages, which is predicted to alter selection for dispersal and resistance. We then test whether bacteria can evolve to escape phages. Combining experimental evolution and mathematical modeling approaches, our findings suggest that phages rarely, if ever, drive the evolution of bacterial escape.

## Materials & Methods

### Bacteria & Phage Strains and Culturing

We used the widely-studied model system of *P. fluorescens* SBW25 and its strictly-lytic phage Phi2 (Brockhurst et al. 2007), which putatively binds the bacterial lipopolysaccharide (LPS) (Scanlan and Buckling 2012). We used variations of King’s B (KB) medium (the standard SBW25 bacterial media, see experimental evolution section below): 10 g/L LP0037 Oxoid Bacteriological Peptone, 15 g/L glycerol, 1.5 g/L potassium phosphate, and 0.6 g/L magnesium sulfate. Amplified phage stocks were created by infecting an actively growing SBW25 liquid culture, incubating while shaking for 24 hours at 28°C, then adding 25 µL chloroform per mL culture and storing at 4°C.

### Measuring Phage & Bacterial Density

Phage and bacterial concentrations were quantified using standard plating and optical density measurements. Phages were quantified in plaque-forming-units per mL (PFU/mL) using a plaque assay: phage suspensions and 100 µL of an overnight bacterial culture in the appropriate KB concentration were added to 4 mL of 0.5% agar KB media at 45°C then poured over 1.5% agar KB plates. After overnight incubation at 28°C plaque forming units were counted. Bacteria were quantified by plating serial dilutions onto 1.5% agar KB plates and counting the number of colony-forming-units (CFU). In liquid culture, bacterial density was quantified by measuring the optical density at 600 nanometers (OD600) with a spectrophotometer, then translating that to CFU/mL using a previously-established standard curve.

### Experimental Evolution

Experimental evolution was conducted in plates containing media with 3 g/L agar. Approximately 22 hours before inoculation or passage, 25 mL of media was poured into each plate with an internal diameter of 90 mm. For the global parasite treatment, media was cooled to 50°C and Phi2 was added before pouring to reach a final phage concentration of 10^6^ PFU/mL. For the control and local parasite treatments, no phage was added before pouring.

Initial inoculations for timepoint 0 were taken from an exponentially growing culture of ancestral *P. fluorescens* that had been shaken at 250 rpm. For the control and global treatments, 5 µL of a bacterial suspension with 5 x 10^7^ CFU/mL was spotted onto the center of the plate. For the local treatment, phages were added to the bacteria to achieve 5 µL of a mixture containing 5 x 10^7^ CFU/mL bacteria and 5 x 10^6^ PFU/mL phages, then spotted onto the center of the plate. These concentrations were chosen so that the initial inoculum droplet of bacteria would be exposed to approximately the same number of phages between the local and global treatments.

After bacteria are inoculated in the center of the plate, nutrient consumption creates a spatial gradient in chemoattractants. This gradient induces bacterial dispersal from the center of the plate towards the periphery, primarily via flagella-driven swimming. Thus, population-level dispersal is the result of both bacterial growth and motility (Fraebel et al. 2017). In these conditions, phages like Phi2 only passively diffuse (Sampedro et al. 2015).

After initial inoculation, populations were then propagated to a new set of plates daily for 14 transfers, while re-creating the same phage spatial distribution each time. After ∼24 hours incubation, a 300 µL pipette tip set to 5 µL was stabbed into the agar immediately past the visible cell front and drawn up slowly, to ensure no bubbles entered the tip. This 5 µL sample was then spotted onto the center of a new plate. For the global treatment, plates were poured in the same manner as initial inoculations, incorporating phages throughout the media. For the local treatment, 2.5 µL of a phage stock with 10^7^ PFU/mL was added onto the 5 µL droplet on the new plate, yielding the same area-density as the initial inoculations. For the control treatment, no phages were added. After 14 transfers, evolved isolates were taken by spreading diluted suspensions of the 5 µL sample onto 1.5% agar KB plates and incubating for 48 hours.

This experiment, with five replicate populations in each of the three parasite distribution treatments (Fig 1), was carried out twice under different media and incubation conditions. In the first experiment, bacteria and phages were incubated at 30°C; in the second experiment, bacteria and phages were incubated at 29°C. Previous work has shown that Phi2 does not grow at 30°C but can grow at 29°C (Padfield et al. 2019). We confirmed this pattern by observing that Phi2 formed plaques on ancestral bacteria at 29°C but not at 30°C. The number of generations was estimated by dividing the total duration of the experiment by the average ancestral growth rate (Fig S1) from each experiment.

**Figure 1.**
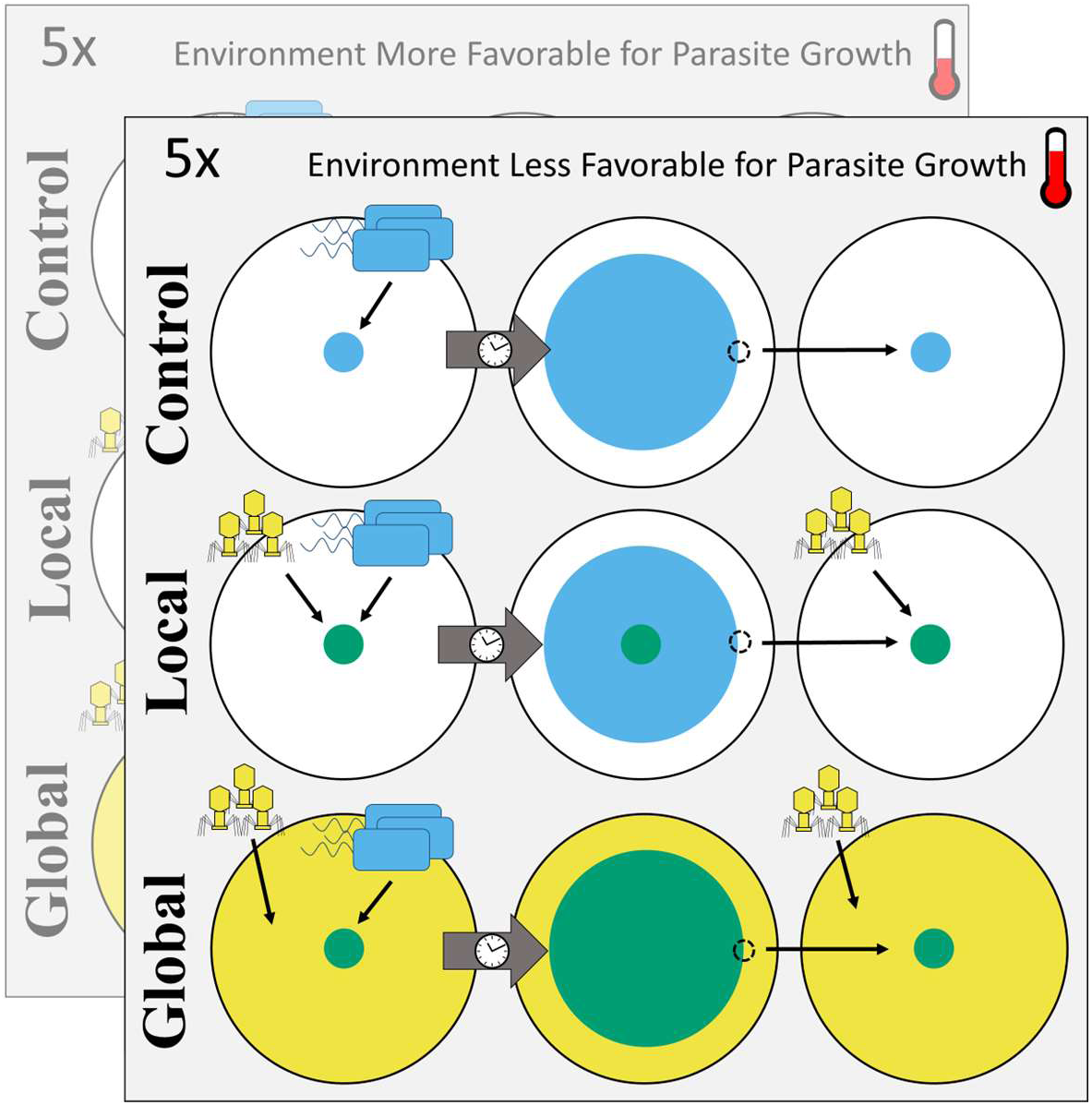
Experimental evolution design. Bacteria were inoculated into the center of an agar plate containing one of three treatments: phages excluded (control), phages only in the center of the plate (local), or phages throughout the plate (global). After 24 hours of incubation, bacteria (blue) have actively dispersed outwards, and a sample was taken from the cell front and transferred to the center of a new plate containing phages in the same distribution. Five evolutionary replicates were carried out for each treatment. This entire experiment was run twice, once under conditions more favorable for phage growth, and once under conditions less favorable for phage growth.

Nutrient conditions also differed between the two experiments. Bacteria growing in media with 3 g/L agar can disperse via swarming or swimming motility (Kohler et al. 2000; Harshey 2003). For experimental tractability, nutrient concentrations in the media were altered to prevent the formation of swarms. At each temperature, we tested KB formulations with 3 g/L agar, either 25%, 50%, 75%, or 100% concentrations of glycerol and peptone (the carbon and nitrogen sources), and 50% concentrations of potassium phosphate and magnesium sulfate. For each experiment, we then selected the highest-nutrient formulation that did not induce bacterial swarming. For the first experiment, in the conditions relatively less favorable for phage growth, the chosen media was “50%/50% KB”; for the second experiment, relatively more favorable for phage growth, the chosen media was “25%/50% KB”. Note that, because the two experiments were not conducted concurrently, and because of the media and temperature differences between the two experiments, comparisons should only be made within each experiment.

Experimental evolution populations were blocked such that each block contained one population in each of the control, local, and global treatments. In some cases, plates showed no growth after 24 hours of incubation. In these cases, all three populations in the block were re-inoculated from the previous transfer, which had been stored at 4°C. This occurred seven times (Table S1), predominantly in the first two transfers, when phage killing was most frequent. Additionally, one global and one local population, both in the experiment with less favorable conditions for phage growth, became contaminated during the experiment and were excluded from analysis.

### Quantifying Bacterial Dispersal/Growth in Soft Agar

Bacterial growth during experimental evolution and dispersal (sometimes called ‘migration rate’ in the literature) of evolved bacterial isolates were measured by photographing plates at 300 dpi. For experimental evolution, plates were photographed immediately before passaging. For evolved isolates, 5 µL of a bacterial suspension at 5x10^7^ CFU/mL was spotted onto the center of a 25mL 0.3% agar plate of the appropriate media, then incubated for approximately 24 hours before photographing. To reduce batch effects, each block was poured from the same bottle of media the day before inoculation, although the drying time was not precisely standardized. After photographing, the area of the circular bacterial growth in these images was measured manually using Fiji (Schindelin et al. 2012). This area was normalized for the exact time of incubation under the assumption that radius increases linearly with time (Croze et al. 2011). For evolved isolates, the batch-corrected dispersal rate is reported.

### Quantifying Bacterial Resistance to Phage

Bacterial resistance to phage was quantified by tittering a stock of the phage on lawns of the ancestral bacteria and evolved isolates, then reporting the batch-corrected number of plaques observed on each isolate.

### Statistical Analysis

All analyses and visualization were carried out in R 4.2.2 (R Core Team 2022) and RStudio 2022.12.0+353 using packages reshape (Wickham 2007), tidyr (Wickham et al. 2023*b*), dplyr (Wickham et al. 2023*a*), ggplot2 (Wickham 2016), ggh4x (Brand 2022), lme4 (Bates et al. 2015), pbkrtest (Halekoh and Højsgaard 2014), gcplyr (Blazanin 2023), cowplot (Wilke 2020), ggrepel (Slowikowski 2022), ggtext (Wilke and Wiernik 2022), magrittr (Bache and Wickham 2022), and purr (Wickham and Henry 2023), along with the Okabe and Ito colorblind-friendly palette (Okabe and Ito 2008). Any other statistical functions, like prcomp, were built-in to R. See https://github.com/mikeblazanin/trav-phage for all analysis code.

### Simulations

To simulate bacterial dispersal with different parasite spatial distributions, we modified the widely-used Patlak-Keller-Segel model of bacterial chemotaxis and growth (Patlak 1953; Keller and Segel 1970) to include phage populations (Eqs. 3 – 6). These models have been previously validated to recreate *in vitro* bacterial population behavior with great precision (Cremer et al. 2019; Li et al. 2020; Ping et al. 2020; Mattingly and Emonet 2022).

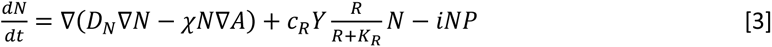

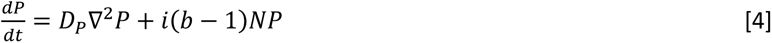

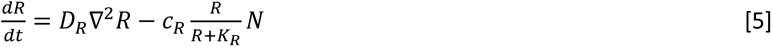

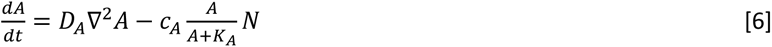

In this model we track the population density of hosts (N), parasites (P), resources (R), and attractants (A) through both space and time, using parameters as defined in Table 1. Simulations were initiated with parasites excluded (Control), gaussian distributed centered on the origin (peak height = 5x10^8^ parasites/*um*, sd = 20*um*) (Local), or uniformly distributed (2.5 parasites/*um*) (Global). Across all treatments, hosts were initiated gaussian distributed centered on the origin (peak height 10^10^ cells/*um*, sd = 20 *um*), resources were initiated uniformly at 50 mM, and attractants were initiated uniformly at 2 mM.

**Table 1.** Parameters and values used in simulations.

| Parameter | Description | Resident Value |
| --- | --- | --- |
| $D_N$ | Diffusion coefficient for bacteria | $50 \mu\text{m}^2/\text{s}$ |
| $D_P$ | Diffusion coefficient for phage | $0 \mu\text{m}^2/\text{s}$ |
| $D_R$ | Diffusion coefficient for resource | $800 \mu\text{m}^2/\text{s}$ |
| $D_A$ | Diffusion coefficient for attractant | $800 \mu\text{m}^2/\text{s}$ |
| $\chi$ | Chemotactic sensitivity coefficient | $315 \mu\text{m}^2/\text{s}$ |
| $c_R$ | Uptake rate of resource | $2.25 \times 10^{-14}$ mmol/cfu/s |
| $c_A$ | Uptake rate of attractant | $4.17 \times 10^{-16}$ mmol/cfu/s |
| Y | Cell yield | $1.229 \times 10^{10}$ cfu/mmol |
| $K_R$ | Michaelis-Menten constant for resource | $5 \times 10^{-5}$ mmol/mL |
| $K_A$ | Michaelis-Menten constant for attractant | $1 \times 10^{-6}$ mmol/mL |
| i | Infection rate | $1 \times 10^{-12}$ infec/cell/mL/pfu/mL/s |
| b | Number of parasites produced per infection | 30 pfu/infec |

To assess the fitness landscape between resistance (i) and dispersal (χ), growth rate (c_R_), attractant consumption (c_A_), or yield (Y), the host population was split into two equally-sized sub-populations that shared the same initial distribution. One sub-population (the resident) had the parameter values listed in Table 1, while the other (the invader) had one or two parameter values which differed from the resident. See Table S3 and the Supplemental Materials for how parameter values were determined. Following this setup, the simulation was run for 20 simulated hours. At the end of the simulation, the fitness of the invader was calculated as log_10_(invader frequency/resident frequency). All models were implemented in Matlab (Mattingly and Emonet 2022) and all code is available at https://github.com/jeremymoore558/ks_phage. Visualization code is available at https://github.com/mikeblazanin/trav-phage.

## Results

### Experimental Evolution of Bacteria

To test how parasite spatial distribution alters host evolution, we experimentally evolved *P. fluorescens* in three treatments differing in their spatial distribution of parasites (Fig 1). This experiment was run twice, once under incubation and media conditions more favorable for phage growth, and once under conditions less favorable for phage growth. We estimate that approximately 450 generations passed in the experiment less favorable to phage growth, and approximately 535 generations passed in the experiment more favorable to phage growth. After each day, the radius of bacterial growth was measured (Fig 2). Within each experiment, we observed transient suppression of bacterial growth during the early experimental evolution transfers in treatments containing phages (local and global). This pattern is consistent with initially high rates of phage predation on bacteria, which weakens following host evolution.

**Figure 2.**
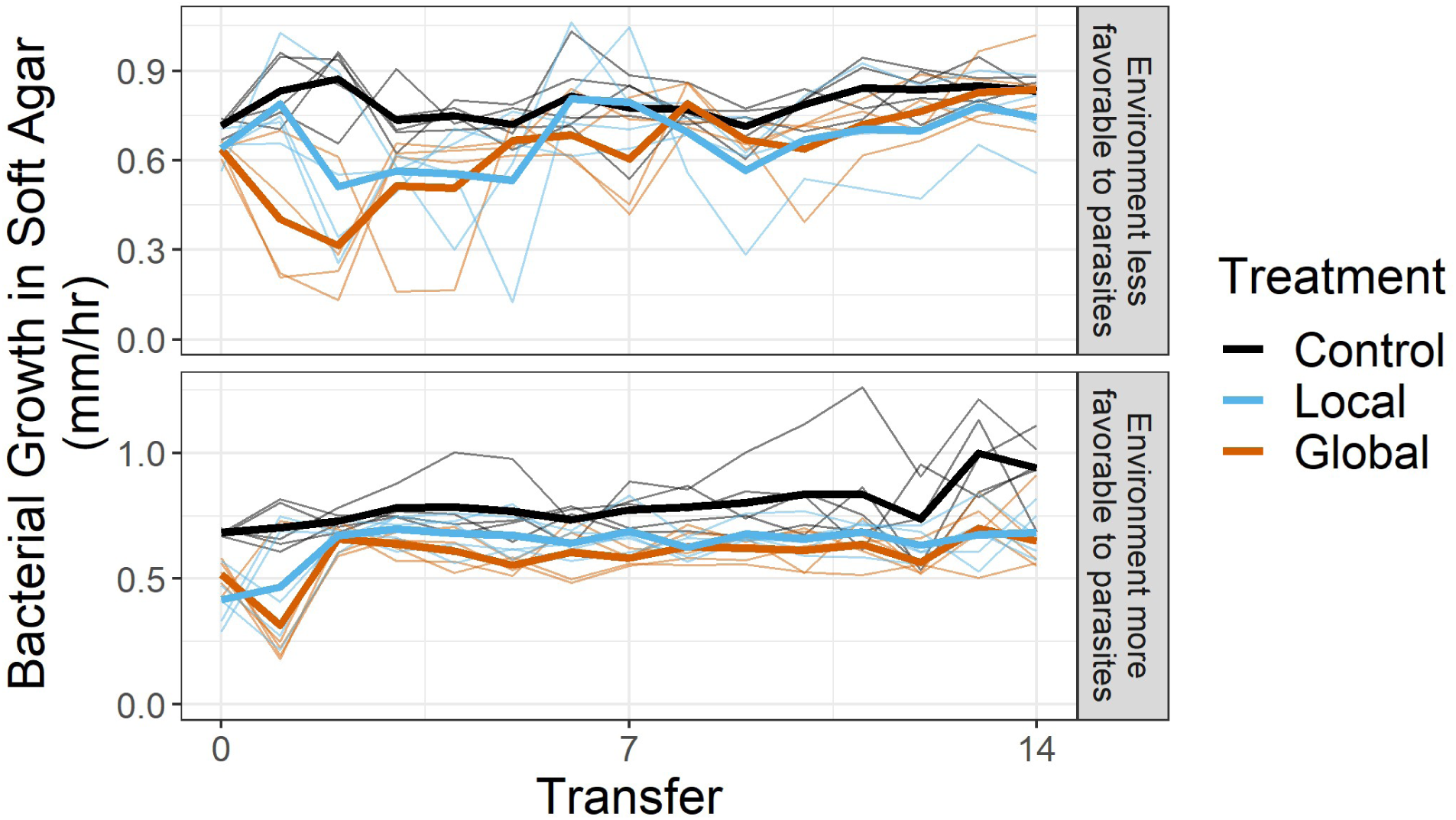
Bacterial populations are initially strongly depleted in the presence of parasites. During experimental evolution, total growth of bacteria was measured at each transfer. Independent populations are plotted as fine lines, and bold lines denote the mean in each treatment. Note the difference in y-axis scales due to incubation and media differences between the experiment with more favorable conditions for phage growth and the experiment with less favorable conditions for phage growth. The less favorable environment local population D and global population C were contaminated during experimental evolution and excluded.

### Isolate Characterization

After experimental evolution, we isolated five evolved bacterial clones from the final timepoint in each replicate population. We then measured how parasite spatial distribution had altered the evolution of two traits: resistance to phage, and dispersal in soft agar.

### Resistance to Phage

We measured the resistance of evolved bacteria to parasite infection (Fig 3). We found that parasite spatial distribution had a substantial effect on resistance evolution. In the experiment less favorable for phage growth, one local population and all global populations evolved increased levels of resistance. In the experiment more favorable for phage growth, all populations in both the local and global treatments evolved increased levels of resistance.

**Figure 3.**
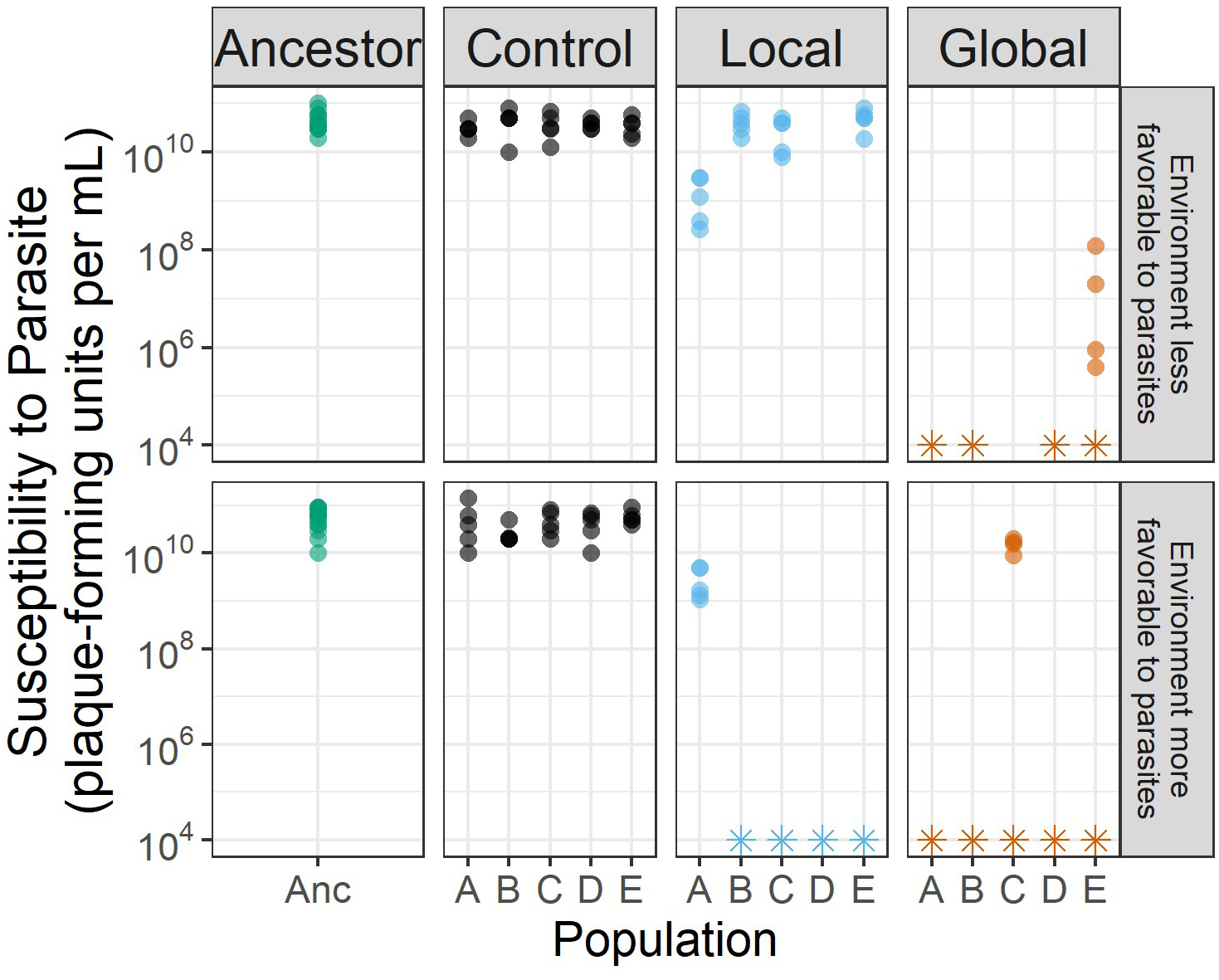
Bacteria evolve reduced parasite susceptibility (resistance) in the presence of globally distributed parasites or in conditions more favorable for parasite growth. Bacterial isolates from the final transfer of experimental evolution were assayed for their susceptibility to phage infection by counting the number of plaques formed by a standard phage stock. Plotted is the number of plaque-forming units, batch-corrected, where isolates with fewer plaque-forming units than control populations could be considered partially resistant. Isolates whose susceptibility fell below the limit of detection (i.e., zero plaque-forming units) are plotted as a star and could be considered fully resistant. Less favorable phage growth local population D and global population C were contaminated during experimental evolution and excluded.

### Dispersal

We measured the dispersal rate of evolved bacteria in soft agar (Figs 4). As expected, evolved isolates tended to have increased dispersal relative to the ancestor, although this effect was largely non-significant (linear mixed-effects model of dispersal with random effects for population and batch and fixed effect for treatment, control-ancestor contrast in more favorable phage condition p = 0.08, all other contrasts with ancestor p > 0.3). However, we failed to find parasite-driven evolution of increased dispersal rate beyond this baseline. Moreover, parasite presence constrained the evolution of increased dispersal in the more favorable phage condition [linear mixed-effects model as above, in less favorable phage condition all contrasts among control, local, and global p > 0.7, in more favorable phage condition control treatment is higher than local (p = 0.06) and higher than global (p = 0.05), while local and global treatments do not differ (p = 0.99)].

**Figure 4.**
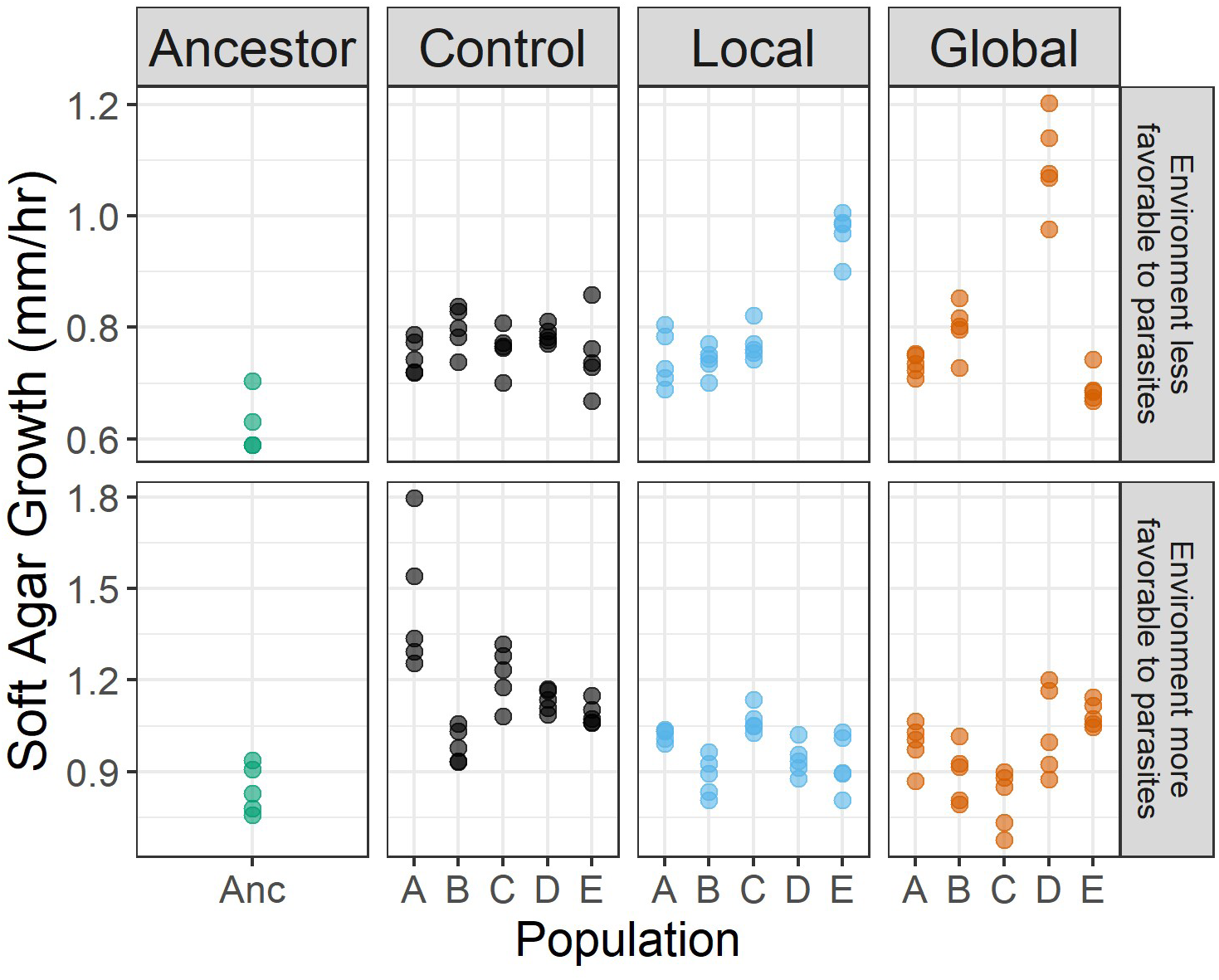
Bacteria evolved increased dispersal, regardless of parasite spatial distribution. Bacterial isolates from the final transfer of experimental evolution were assayed for their dispersal in soft agar in the absence of parasites. Plotted are dispersal rates, batch corrected. Note the difference in y-axis scales due to incubation and media differences between the experiment more favorable for phage growth and the experiment less favorable for phage growth. Less favorable phage growth local population D and global population C were contaminated during experimental evolution and excluded.

### Mathematical Modeling

These results showing the evolution of resistance and not dispersal could be specific to any number of biological particularities of our model system. Unfortunately, establishing the generality of these findings *in vitro* is experimentally intractable (Lustenhouwer et al. 2023). Instead, we leveraged existing mathematical models of bacterial growth and dispersal to simulate bacterial evolution in the presence of different parasite spatial distributions across a wide range of bacterial trait values. Using *in silico* invasion experiments, we uncovered the fitness landscape between dispersal and resistance and how it shifted depending on parasite spatial distribution (Fig 5). As expected, in the absence of parasites, resistance provides no benefits, and dispersal is selected for since it enables faster migration into space with higher resource availability (Mattingly and Emonet 2022). Surprisingly, in the presence of parasites, regardless of their distribution, this pattern is exactly flipped: resistance is selected for, while dispersal provides no benefits. This arises because phage killing reduces bacterial population densities, which weakens the gradient in attractants that induces dispersal, and the gradient in resources that provides benefits to faster dispersers. Moreover, this pattern is robust to the form of the global distribution (Fig S8), and is in sharp contrast to patterns observed with other traits, like growth rate or yield, where parasite presence strengthens selection for resistance without weakening selection for the alternative trait (Figs S9-S11).

**Figure 5.**
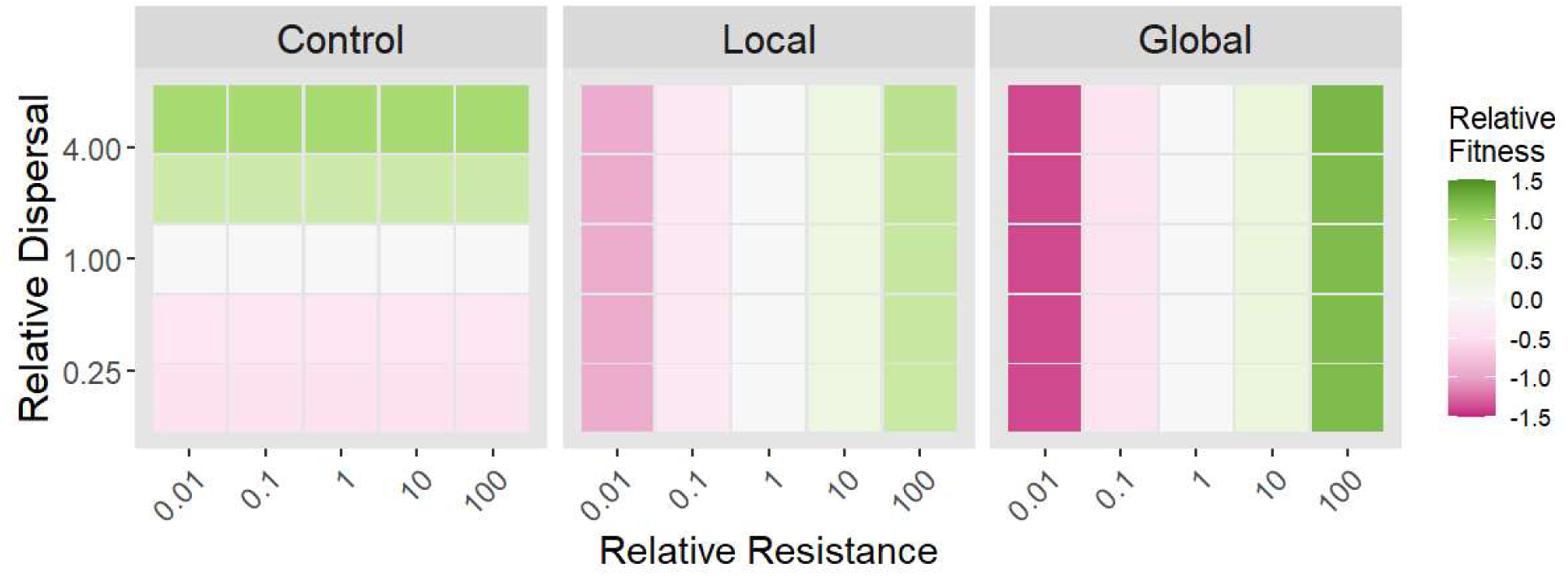
Parasite presence selects for host resistance and eliminates selection for host dispersal. *In silico* invasion experiments were carried out with a bacterial mutant that varied in its traits relative to the resident bacterial population, both competing in the presence of one of three different initial parasite spatial distributions. Results are colored by invader fitness = log_10_(final invader frequency/final resident frequency). Resistance is defined as the reciprocal of infection rate (i), and dispersal is the chemotactic sensitivity parameter (χ).

## Discussion

Here we directly tested whether bacterial dispersal can evolve to escape local parasites. In our *in vitro* experimental evolution, we found that bacterial hosts adapted to parasites not by the evolution of dispersal, but via the evolution of resistance (Figs 3-4). Our *in silico* simulations expanded the generality of our empirical conclusions, showing that parasite presence actually weakens selection for greater dispersal (Fig 5).

Our findings build on past studies of bacterial dispersal in semi-solid environments in the presence of parasitic phages. In particular, our work contradicts the conclusions of two papers whose results implied bacteria could evolve dispersal to escape phages. Li et al. studied the ecological dynamics of *Salmonella enterica* as it dispersed in soft agar in the presence of χ phages (Li et al. 2020). Based on short term experiments, they predicted that over longer evolutionary time scales, bacteria and phages could be in a coevolutionary arms race to evolve faster dispersal. Second, Taylor & Buckling studied the competitive benefits of *Pseudomonas aeruginosa* motility in the presence of different spatial distributions of phage PhiKZ (Taylor and Buckling 2013). They found that motility was beneficial in the presence of phages regardless of the spatial distribution of those phages. Based on those results, they argued that phages have the potential to select for bacterial dispersal. These two ecological studies extrapolated evolutionary outcomes, but those outcomes did not materialize in our direct evolutionary tests, where we find that phages do not select for bacterial dispersal (Figs 4, 5).

In contrast, our findings do align with, and extend, the work of Ping et al., who studied the ecology and evolution of *E. coli* and phage P1 as they disperse in soft agar from a shared inoculation site (Ping et al. 2020). Their experiments and mathematical modeling showed that bacteria face selection for resistance when they spread slower than dispersing phages, and implied that bacteria would not face selection for dispersal when they are already spreading faster than dispersing phages. However, they did not directly test whether phages would alter selection on dispersal, and their modeling assumed a constant cost of phage resistance. In this study, we directly confirmed the implications of their work: phages do *not* select for bacterial dispersal (Figs 4, 5), and resistance is favored over avoidance across the whole fitness landscape (Fig 5).

More broadly, our work contributes to the nascent study of avoidance in host-parasite evolution, and suggests that avoidance via escape may be fundamentally less relevant in microbial systems than in multicellular systems. To our knowledge, this field has nearly-exclusively focused on multicellular host species (Gibson and Amoroso 2022), where avoidance is fairly evolvable (Gibson and Amoroso 2022), and can be a low-cost and effective strategy (Kiesecker et al. 1999; Taylor et al. 2004; Daly and Johnson 2011). Multicellular hosts have been found to avoid parasites by limiting behavior that could expose them to parasites (Stephenson et al. 2018; Paciência et al. 2019), or by dispersing to escape local threats of parasitism (Sloggett and Weisser 2002; Fill et al. 2012; Baines et al. 2020; Brophy and Luong 2021; Zilio et al. 2021). In contrast, microbes have only previously been found to avoid parasites by limiting swarming that could expose them to parasites (Bru et al. 2019), and our work suggests that the evolution of escape may be rare or non-existent in microbes. For instance, microbial dispersal provides little benefit in escaping local parasites (Fig 5), and is substantially less evolvable than resistance (Figs 3, 4), where a single mutation can alter susceptibility by many orders of magnitude (Chan et al. 2016; Burmeister et al. 2020; Kortright et al. 2020). In all, these findings imply that generalized microbial dispersal likely functions primarily as a foraging strategy (Keegstra et al. 2022) and not as a mechanism to escape parasites.

Our results generally follow previously reported trends on the evolution of bacteria under selection by phages. At the beginning of the evolution experiment, we observed decline and subsequent recovery of bacterial population size (Fig 2). Such dynamics are common in phage-bacteria experiments where bacteria are initially susceptible (Koskella et al. 2011). Our finding of phage resistance in the global treatment (Fig 3) also echoes past work where more homogenous environments accelerated the coevolution of bacterial resistance with phages (Brockhurst et al. 2003). Additionally, the maintenance of phage susceptibility in our local phage treatment (Fig 3) is consistent with past work that spatial structure and heterogeneity increases host-parasite coexistence (Brockhurst et al. 2006; Tamar and Kishony 2022).

We were somewhat surprised to find the maintenance of intra-population variation in resistance, especially in bacteria from the global treatment (Fig 3). Foremost, this is surprising because bacteria in the global treatment generally experienced strong selection for parasite resistance, and in other studies of phage selection resistance often rapidly sweeps to fixation (Bohannan and Lenski 2000; Buckling and Rainey 2002). In addition, populations undergoing range expansion are subject to strong drift via ‘allele surfing’ at the expanding frontier (Hallatschek et al. 2007; Excoffier and Ray 2008). Since we sampled the cell front for our experimental transfers, this should further limit the maintenance of diversity. However, several mechanisms could explain the unexpected maintenance of diversity, including phenotypic plasticity or switching (Tzipilevich et al. 2022), balancing or diversifying selection for resistance (Bohannan and Lenski 2000; Boots et al. 2008), or collective migration that maintains diversity (Fu et al. 2018).

There are many avenues for future work opened by our findings. First, we used bacterial and phage strains where there is no *a priori* expectation for linkage between resistance and dispersal traits [Phi2 putatively binds bacterial LPS as a receptor (Scanlan and Buckling 2012)]. However, many phage-bacteria systems do have a mechanistic link between dispersal and resistance. For instance, flagellotropic phages attach to and infect their host cells via the bacterial flagella (Icho and Iino 1978). In these systems, bacterial resistance often evolves by loss or modification of the flagella, creating a tradeoff between motility and resistance. Future work could investigate the evolution of resistance and avoidance in systems like these where the two traits are pleiotropically linked. Additionally, our work focused exclusively on one-sided evolution. We designed our experiments to minimize phage coevolution, since phage dispersal from the inoculation site to the sampling site at the cell front is likely rare (Ping et al. 2020), and because the ancestral phage was reintroduced at each passage. Experiments can be designed to explicitly incorporate coevolution in a spatially structured environment (Tamar and Kishony 2022), but whether coevolution alters the evolution of resistance and avoidance remains to be tested.

Bacteria have abundant strategies to resist phage infection (Hampton et al. 2020), including the capability to activate defenses when they detect infections in nearby neighbors (de Mattos et al. 2022), as well as complex abilities to sense and physically navigate their environment (Keegstra et al. 2022; Wadhwa and Berg 2022). Indeed, both eukaryotes and bacteria have been found to avoid parasite exposure (Bru et al. 2019; Gibson and Amoroso 2022). However, despite the widespread existence of escape from parasites in mobile eukaryotes (Gibson and Amoroso 2022), our experiments and, especially, theoretical results suggest that bacterial escape from parasites may only rarely, if ever, be selected for.

## Supplementary Materials

### Repeated passages

**Table S1.** Repeated Passages Due to Sampling. During experimental evolution seven transfers failed to show growth in at least one treatment after 24 hours, likely due to sampling effects of the low-density cell front. In these instances, a new set of plates was re-inoculated from the previous plates, which had been stored at 4C.

| Experiment | Replicate Population | Transfer |
| --- | --- | --- |
| Less favorable for phage growth | A | 2 |
| Less favorable for phage growth | B | 2 |
| Less favorable for phage growth | B | 6 |
| Less favorable for phage growth | B | 7 |
| More favorable for phage growth | C | 1 |
| More favorable for phage growth | D | 1 |
| More favorable for phage growth | D | 2 |

### Evolved growth

During experimental evolution, bacteria may have adapted their growth characteristics and life history traits to the different treatments. Alternatively, these traits may have evolved following trade-offs with other traits that were under selection, like resistance to phages or dispersal in soft agar. To measure growth characteristics, we collected growth curves of evolved isolates from the final timepoint in liquid media. Each isolate was grown in their experimental evolution environment (“original”), as well as an environment with all nutrients doubled in concentration (“rich”). This rich media was included to possibly reveal cases when bacteria had evolved growth costs due to trade-offs with dispersal or resistance that had then been ameliorated by compensatory mutations. By doubling the nutrients, we hypothesized this may reveal any costs of evolution hidden by compensatory mutations.

Growth curves were computationally characterized, with computational findings manually validated, as described below. In particular, we identified the following traits:

- Maximum per-capita growth rate (r)
- Lag time (time when maximum per-capita growth first exceeded 0.5/hr, or 0.4/hr when 0.5 was never reached)
- Time when a diauxic shift occurred, if any (k_t_)
- Density when a diauxic shift occurred, if any (k)
- Deceleration parameter as bacterial density approaches diauxic shift (v)

There was no significant effect of parasite distribution or evolved resistance on maximum per-capita growth rate (Fig S1, Table S2), and there were no notable effects in other growth characteristics (Figs S2, S3, S4, S5).

### Methods for Quantifying Bacterial Growth in Liquid

To quantify bacterial growth in liquid, 4 µL of an overnight bacterial culture was added, in duplicate wells, to 146 µL of media in each well of a 96-well plate. Media included both the media experimental evolution was carried out in (“original”), and media with the concentrations of all components doubled from that (“rich”). This plate was shaken and incubated at the appropriate temperature overnight, with the OD600 read every 15 or 30 minutes.

Then, growth curve data was computationally characterized using gcplyr (Blazanin 2023) and fitted to determine isolates’ lag time, maximum per-capita growth rate (r), and density and time when a diauxic shift was observed (k, k_t_). The first half hour of data points were noisy, so they were excluded. Then, data was smoothed using a moving median of 3 points, followed by LOcally Etimated Scatterplot Smoothing (LOESS) with a span of either 1 hour or 6.94 hours. We identified the exponential start of logistic growth as the first point when the per-capita growth rate of the of the 1-hour LOESS-smoothed data exceeded 0.5/hour, or 0.4/hour when 0.5 was never reached. We identified the end of logistic growth as the first local minima in the derivative of the 6.94-hour LOESS-smoothed data. These criteria were manually verified as yielding time spans where the data reflected logistic growth, excluding any acclimation period at the beginning of the data, and any post-diauxic shift growth at the end of the data.

We then fit the data in the logistic growth window with a differential equation for a modified form of logistic growth (Baranyi and Roberts 1994; Ram et al. 2019) (Equation 1). This form includes a deceleration parameter, v, to better fit bacterial density as it approaches carrying capacity. Here, N(t) denotes the density of bacteria at time t, r is the maximum per-capita growth rate, k is the carrying capacity, and v is the deceleration curve parameter.

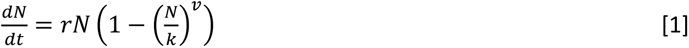

This differential equation has a closed-form solution (Ram et al. 2019) (Equation 2).

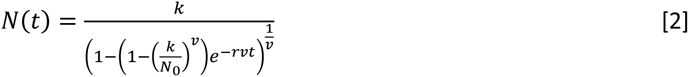

Equation 2 was fitted to the raw unsmoothed data points in the logistic growth window. We then manually evaluated all wells with high fit error, excluding wells where the fit was poor. We extracted fitted maximum per-capita growth rates (r) and excluded any isolates where the standard deviation of r between technical replicates was greater than 0.12/hr (for less favorable phage growth condition) or 0.14/hr (for more favorable phage growth condition). These thresholds were chosen heuristically to exclude wells with large variation between replicates.

### Relationships among evolved traits

To identify relationships among evolved traits in bacterial isolates, including dispersal, resistance to phages, and growth characteristics, we merged all three datasets characterizing the ancestral and evolved isolates. We then excluded any isolates with missing data. We carried out checks for univariate normalcy of all variables, and transformed some variables for improved normalcy: resistance was -log_10_(Efficiency of Plaquing), and diauxic shift density (k) was log_10_(k). Data from the relatively slower growing phage condition-evolved bacteria was multivariate normal, while data from the relatively faster growing phage condition-evolved bacteria was not multivariate normal. Data was scaled and centered and principal component analyses were run.

Generally, multivariate analyses reveal that growth traits were highly correlated between the original and rich medias within a condition (relatively slower growing phage or relatively faster growing phage) (Fig S6). In contrast, growth traits tended to have inconsistent relationships across the two conditions. We observe that global treatment isolates have most diverged from the ancestor in relatively slower growing phage conditions, while control treatment populations have most diverged from the ancestor in relatively faster growing phage conditions (Fig S7).

Previous work has also commonly found relationships between evolved bacterial traits, something we failed to consistently find in our data (Figs S6, S7). For instance, past work has documented tradeoffs between phage resistance and competitive ability (Brockhurst et al. 2004), dispersal and growth rate (Fraebel et al. 2017), or found that evolution with phages can constrain adaptation to the abiotic environment (Koskella et al. 2011; Scanlan et al. 2015). That said, some studies have also failed to find relationships between phage resistance, bacterial motility, and growth, as we did (Koskella et al. 2011). Further work is needed to disentangle the factors driving these differences between studies.

### Simulations parameterization

D_P_ was set to zero because phage particles lack any active motility, and passive diffusion of viral particles through agar gel is very slow relative to the diffusion of nutrients and attractants, or diffusion and migration of bacteria (Ping et al. 2020).

Y was approximated from the observed growth curve data (Fig S3), where bacterial cultures reached carrying capacity on the primary resource around 10^9^ cfu/mL (in the richer media of the slower growing phage condition). Since that media contains 7.5 g/L glycerol (the primary carbon source, molar mass = 92.09 g/mol), this implies a yield of approximately 1.229 × 10^10^ cfu/mmol of glycerol. The range of invader values was set to include and exceed the variation we observed in carrying capacities in our experimental data.

c_R_ was approximated from the observed growth curve data (Fig S1), where bacteria doubled slightly slower than once per hour. For simplicity, we approximated the maximum bacterial growth rate (r) as 1 /hour. Given the known cell yield and primary carbon source concentration (above), this implies a resource uptake rate 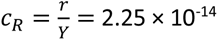 mmol/cfu/s. The range of invader values was set to include and exceed the variation we observed in maximum growth rate in our experimental data.

### Simulation results

As described in the main text, we leveraged existing mathematical models of bacterial growth and dispersal to simulate bacterial evolution in the presence of different parasite spatial distributions across a wide range of bacterial trait values. Using *in silico* invasion experiments, we uncovered the fitness landscape between resistance and other bacterial traits. In the main text we report the fitness landscape between resistance and dispersal with three initial parasite distributions: none, gaussian-distributed (local), and uniformly-distributed (global) (Fig 6). Here, we include an additional “global” parasite distribution that uses a wider gaussian distribution, controlling for possible effects of gaussian vs uniform initial parasite distributions (Fig S8). We also report fitness landscapes between resistance and growth rate (Fig S9), yield (Fig S10), or attractant consumption (Fig S11), as well as fitness landscapes between burst size and resistance (Fig S12), dispersal (Fig S13), growth rate (Fig S14), yield (Fig S15), or attractant consumption (Fig S16).

These fitness landscapes reveal that, in the absence of parasites, increased dispersal, growth rate, and yield are adaptive, while (as expected) resistance is neutral. In the presence of parasites, resistance becomes adaptive. In landscapes of resistance with growth rate and yield, this has the effect of making both resistance and the other trait adaptive. However, strikingly, in the landscape of resistance and dispersal, dispersal loses its fitness benefits, suggesting that the unique dynamics of resistance and dispersal evolution eliminate the benefits of dispersal in the presence of parasites. These patterns only become more pronounced when comparing local to global to global (gaussian) initial parasite distributions. In contrast to resistance, burst size provides little fitness benefit, regardless of parasite spatial distribution.

**Figure S1.**
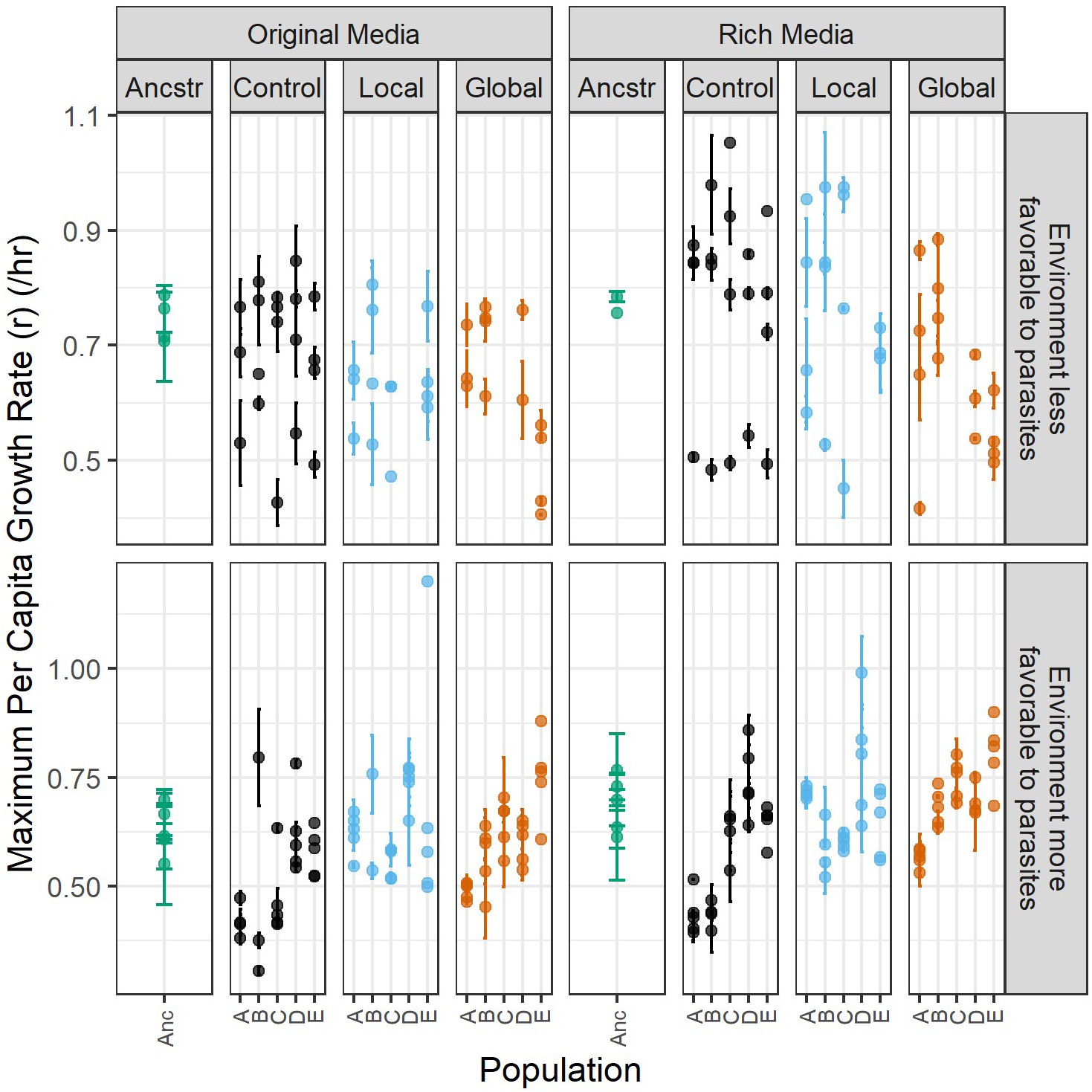
Evolved bacterial growth rates show no effects of parasite distribution. Bacterial isolates from the final transfer of experimental evolution were grown in liquid media containing the same (original) or double (rich) the nutrients as experienced during experimental evolution, quantifying their maximum per-capita growth rate. Each point represents the mean of two replicate wells containing the same isolate, with isolates from the same population stacked vertically, and error bars denoting the standard deviation between replicate wells. Note the difference in y-axis scales due to incubation and media differences between the experiment more favorable for phage growth and the experiment less favorable for phage growth. Less favorable phage growth local population D and global population C were contaminated during experimental evolution and excluded.

**Figure S2.**
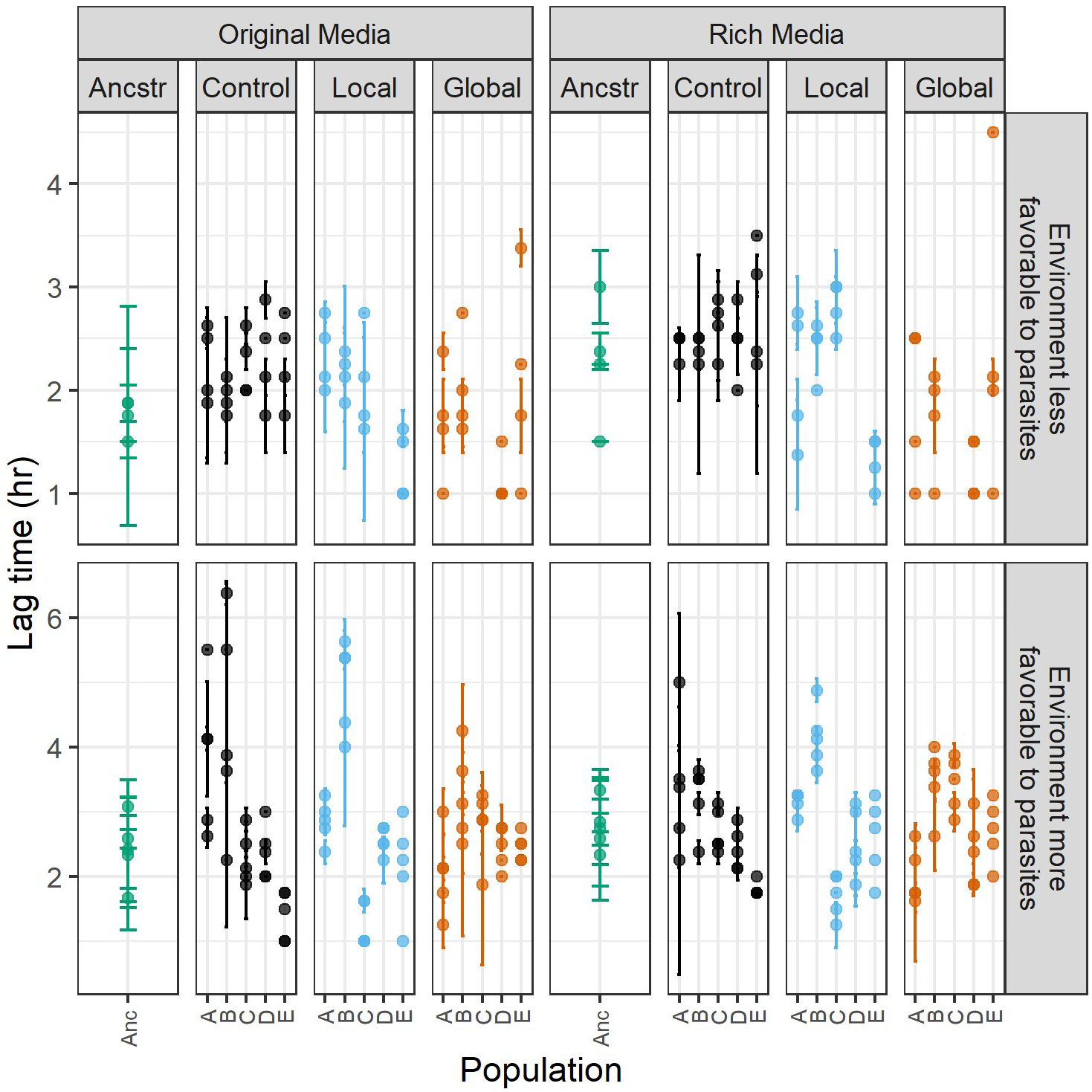
Evolution of lag time. Growth curves of ancestral and evolved bacterial isolates were collected in liquid media containing the same (original) or double (rich) the nutrients as experienced during experimental evolution. Each point represents the mean of two replicate wells containing the same isolate, with isolates from the same population stacked vertically, and error bars denoting the standard deviation between replicate wells. Note the difference in y-axis scales due to incubation and media differences between the experiment more favorable for phage growth and the experiment less favorable for phage growth. Less favorable phage growth local population D and global population C were contaminated during experimental evolution and excluded.

**Figure S3.**
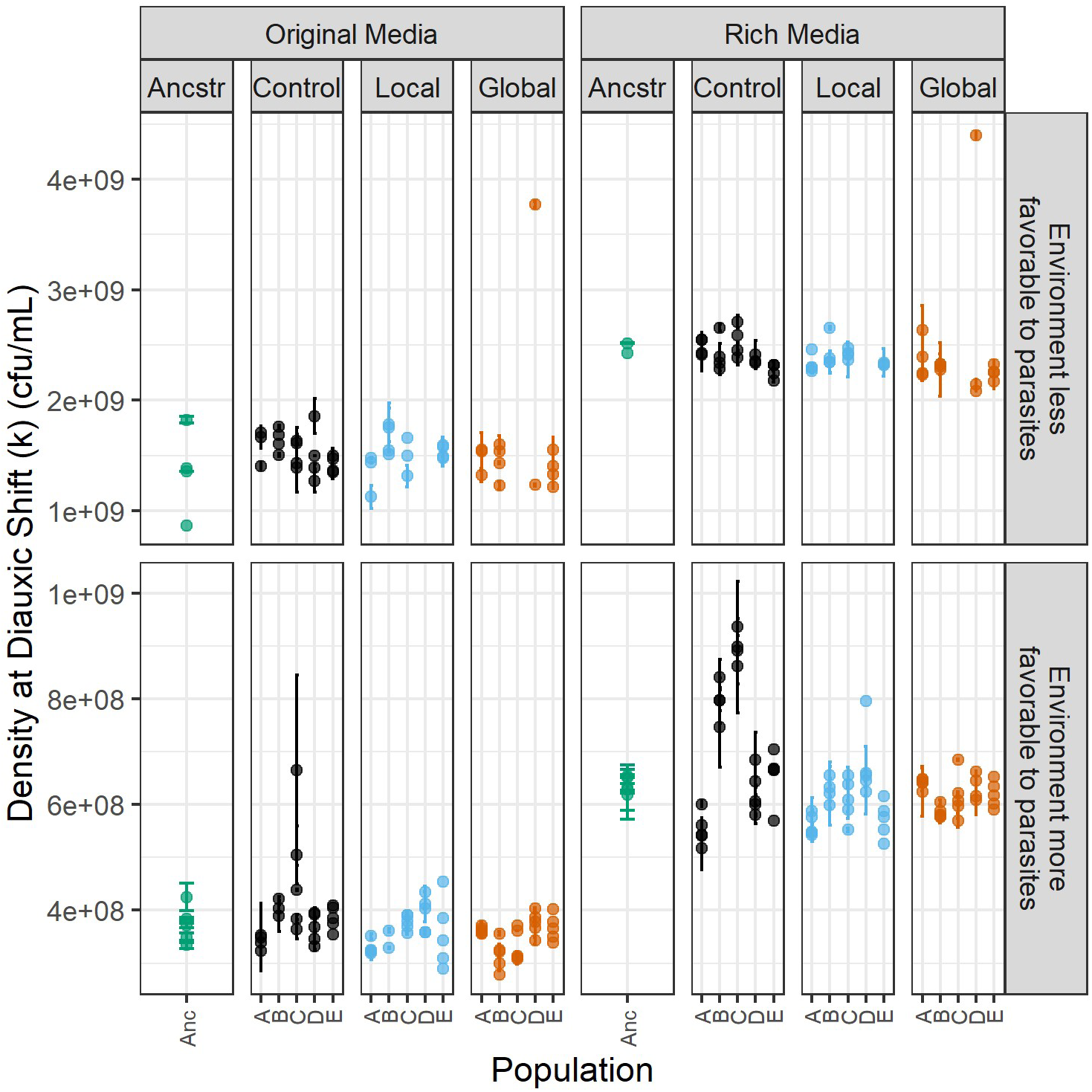
Evolution of diauxic shift density. Growth curves of ancestral and evolved bacterial isolates were collected in liquid media containing the same (original) or double (rich) the nutrients as experienced during experimental evolution. Each point represents the mean of two replicate wells containing the same isolate, with isolates from the same population stacked vertically, and error bars denoting the standard deviation between replicate wells. Points that are outliers with high diauxic shift densities were curves where the bacteria did not undergo a computationally identifiable diauxic shift. Note the difference in y-axis scales due to incubation and media differences between the experiment more favorable for phage growth and the experiment less favorable for phage growth. Less favorable phage growth local population D and global population C were contaminated during experimental evolution and excluded.

**Figure S4.**
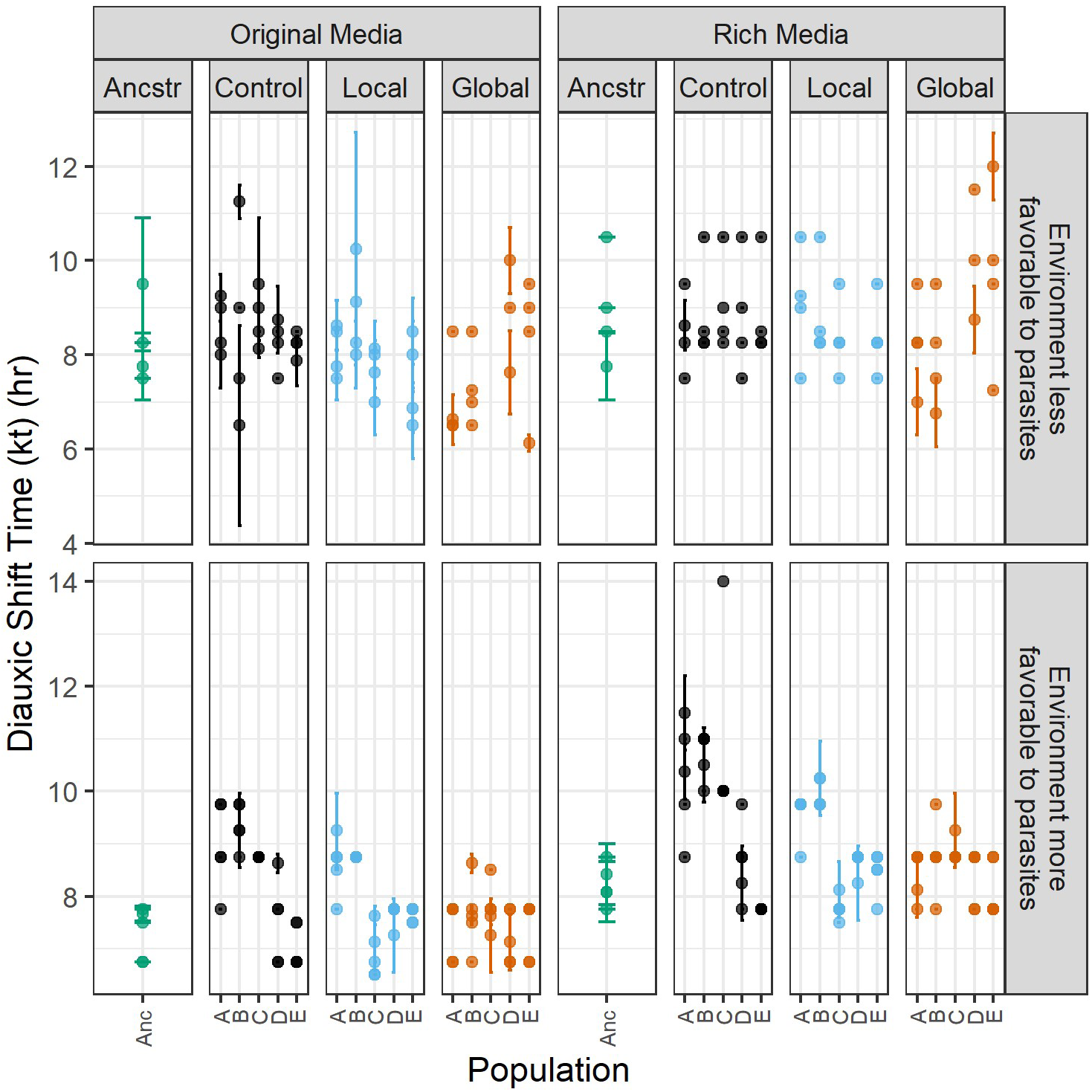
Evolution of diauxic shift timing. Growth curves of ancestral and evolved bacterial isolates were collected in liquid media containing the same (original) or double (rich) the nutrients as experienced during experimental evolution. Each point represents the mean of two replicate wells containing the same isolate, with isolates from the same population stacked vertically, and error bars denoting the standard deviation between replicate wells. Note the difference in y-axis scales due to incubation and media differences between the experiment more favorable for phage growth and the experiment less favorable for phage growth. Less favorable phage growth local population D and global population C were contaminated during experimental evolution and excluded.

**Figure S5.**
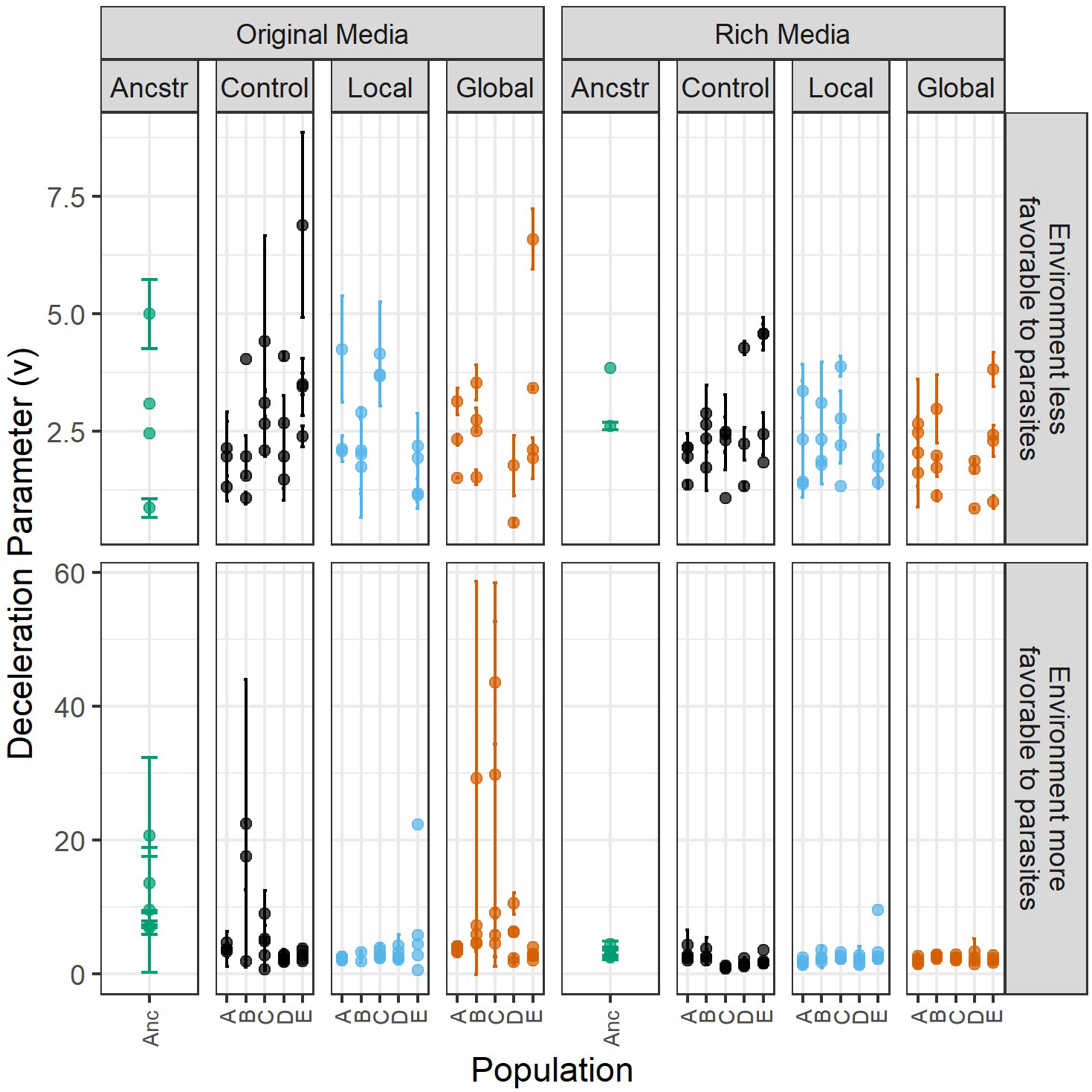
Evolution of deceleration parameter. Growth curves of ancestral and evolved bacterial isolates were collected in liquid media containing the same (original) or double (rich) the nutrients as experienced during experimental evolution. Each point represents the mean of two replicate wells containing the same isolate, with isolates from the same population stacked vertically, and error bars denoting the standard deviation between replicate wells. Note the difference in y-axis scales due to incubation and media differences between the experiment more favorable for phage growth and the experiment less favorable for phage growth. Less favorable phage growth local population D and global population C were contaminated during experimental evolution and excluded.

**Figure S6.**
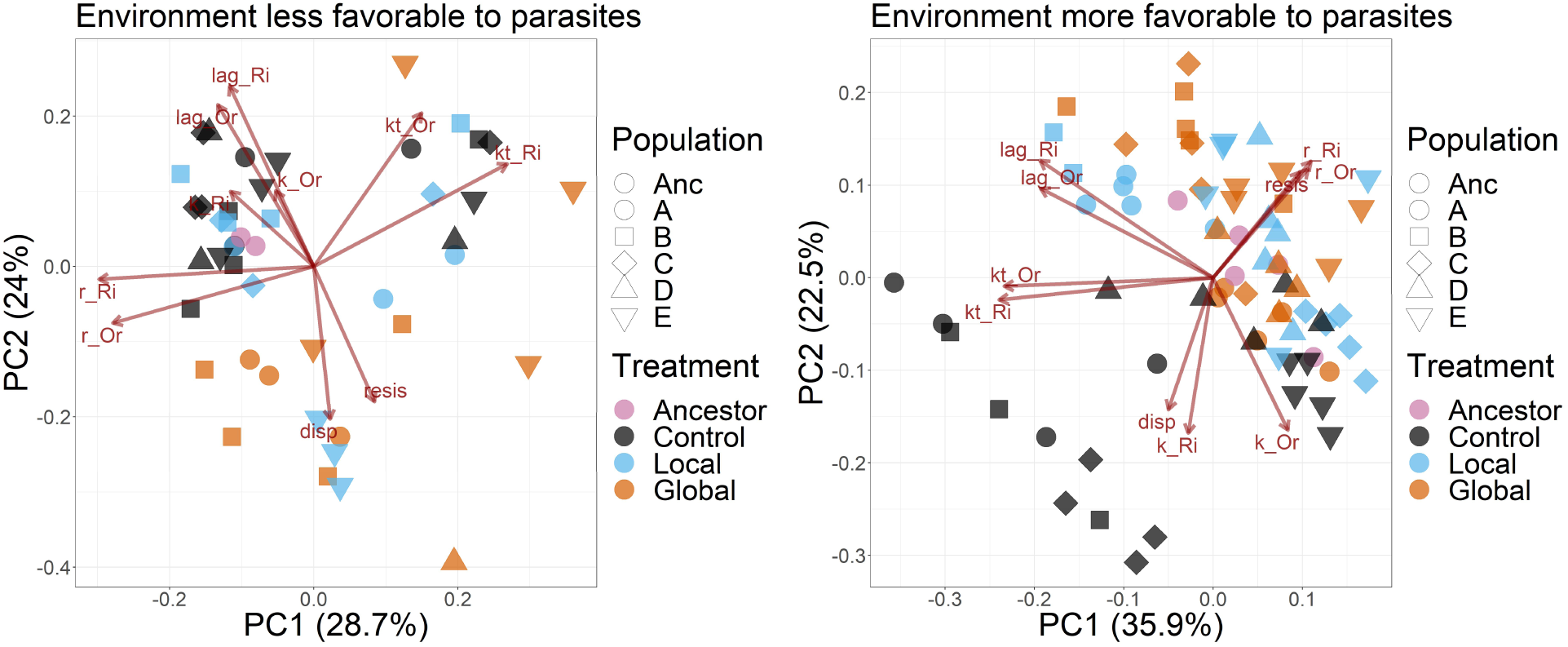
Multivariate evolution of bacterial traits. Data from growth curves, dispersal, and resistance assays were combined, normalized, and principal component analyses were run. Shown is a correlation biplot for each of the two conditions. Each point corresponds to a single isolate. Resis is -log_10_(Efficiency of Plaquing) and disp is dispersal rate. Growth characteristics of lag time (lag), maximum per-capita growth rate (r), the density at diauxic shift (k), and the time of diauxic shift (kt) were measured in media containing the original concentration of nutrients as used in experimental evolution (Or), and rich media containing double the nutrient concentration (Ri). Note that there are incubation and media differences between the environment more favorable to phage and the environment less favorable to phage.

**Figure S7.**
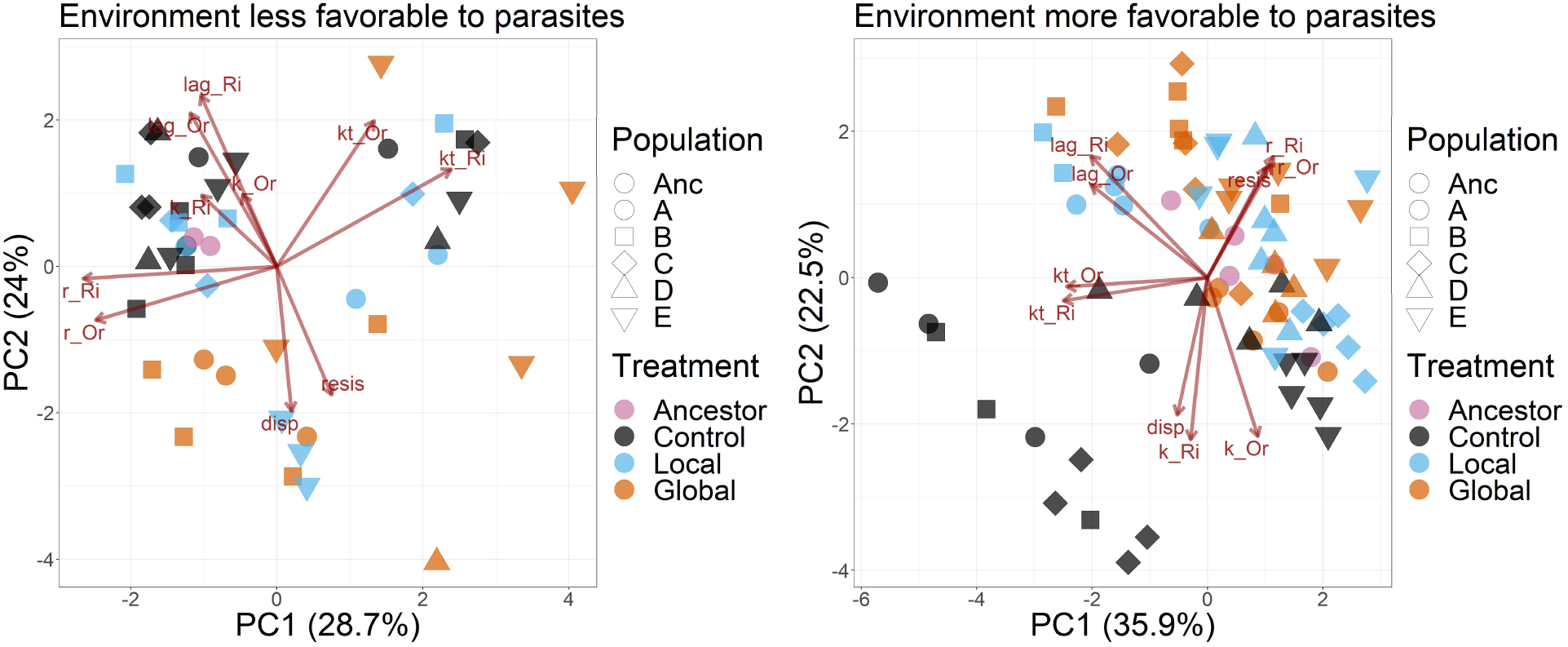
Multivariate evolution of bacterial traits. Data from growth curves, dispersal, and resistance assays were combined, normalized, and principal component analyses were run. Shown is a distance biplot for each of the two conditions. Each point corresponds to a single isolate. Resis is -log_10_(Efficiency of Plaquing) and disp is dispersal rate. Growth characteristics of lag time (lag), maximum per-capita growth rate (r), the density at diauxic shift (k), and the time of diauxic shift (kt) were measured in media containing the original concentration of nutrients as used in experimental evolution (Or), and rich media containing double the nutrient concentration (Ri). Note that there are incubation and media differences between the environment more favorable to phage and the environment less favorable to phage.

**Fig S8.**
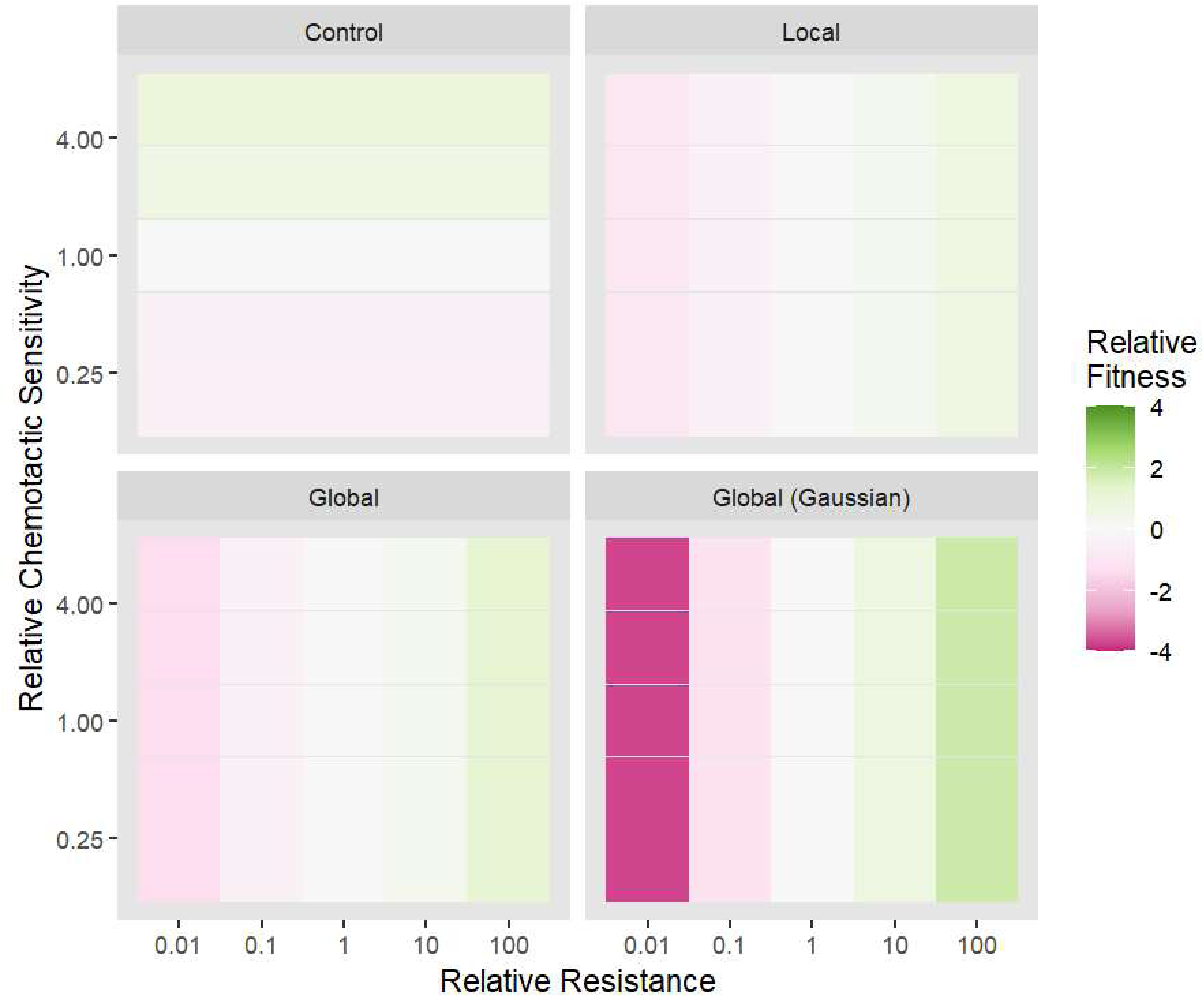
Parasite presence selects for host resistance and eliminates selection for host dispersal. *In silico* invasion experiments were carried out with a bacterial mutant that varied in its traits relative to the resident bacterial population, both competing in the presence of one of four different initial parasite spatial distributions. Results are colored by invader fitness = log_10_(final invader frequency/final resident frequency). Resistance is defined as the reciprocal of infection rate (i).

**Fig S9.**
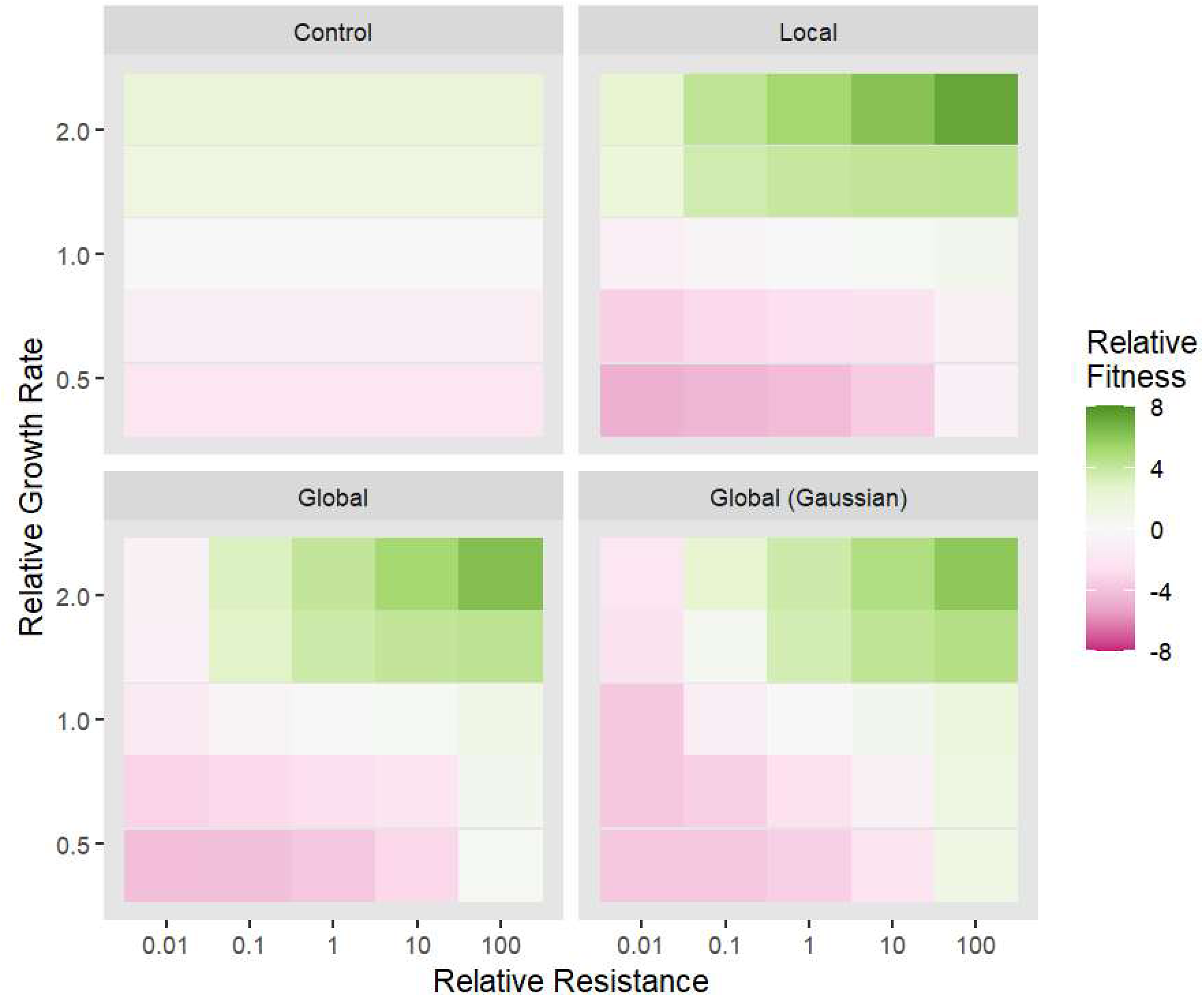
Parasite presence selects for host resistance and maintains selection for host growth rate. *In silico* invasion experiments were carried out with a bacterial mutant that varied in its traits relative to the resident bacterial population, both competing in the presence of one of four different initial parasite spatial distributions. Results are colored by invader fitness = log_10_(final invader frequency/final resident frequency). Resistance is defined as the reciprocal of infection rate (i).

**Fig S10.**
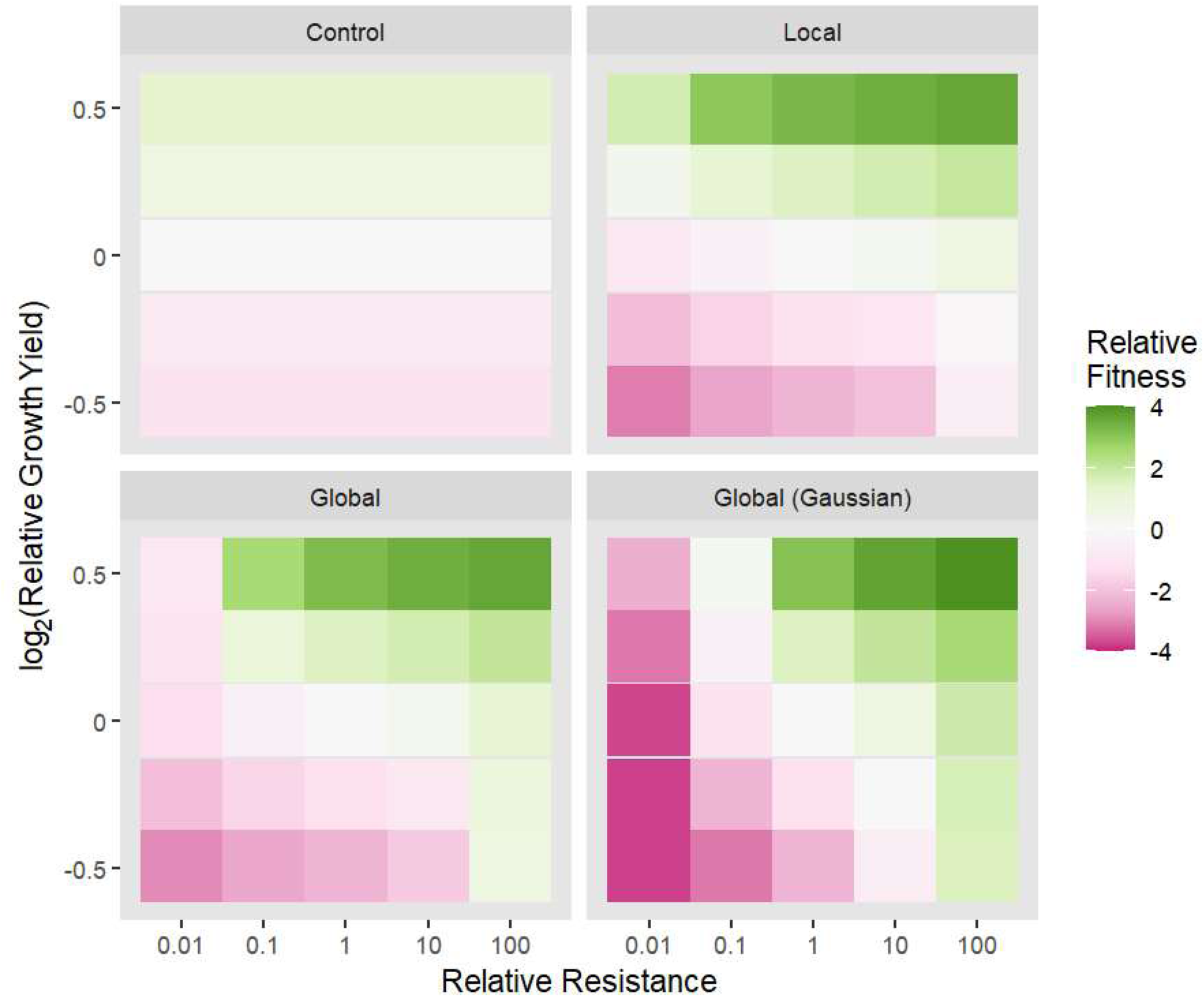
Parasite presence selects for host resistance and maintains selection for host growth yield. *In silico* invasion experiments were carried out with a bacterial mutant that varied in its traits relative to the resident bacterial population, both competing in the presence of one of four different initial parasite spatial distributions. Results are colored by invader fitness = log_10_(final invader frequency/final resident frequency). Resistance is defined as the reciprocal of infection rate (i).

**Fig S11.**
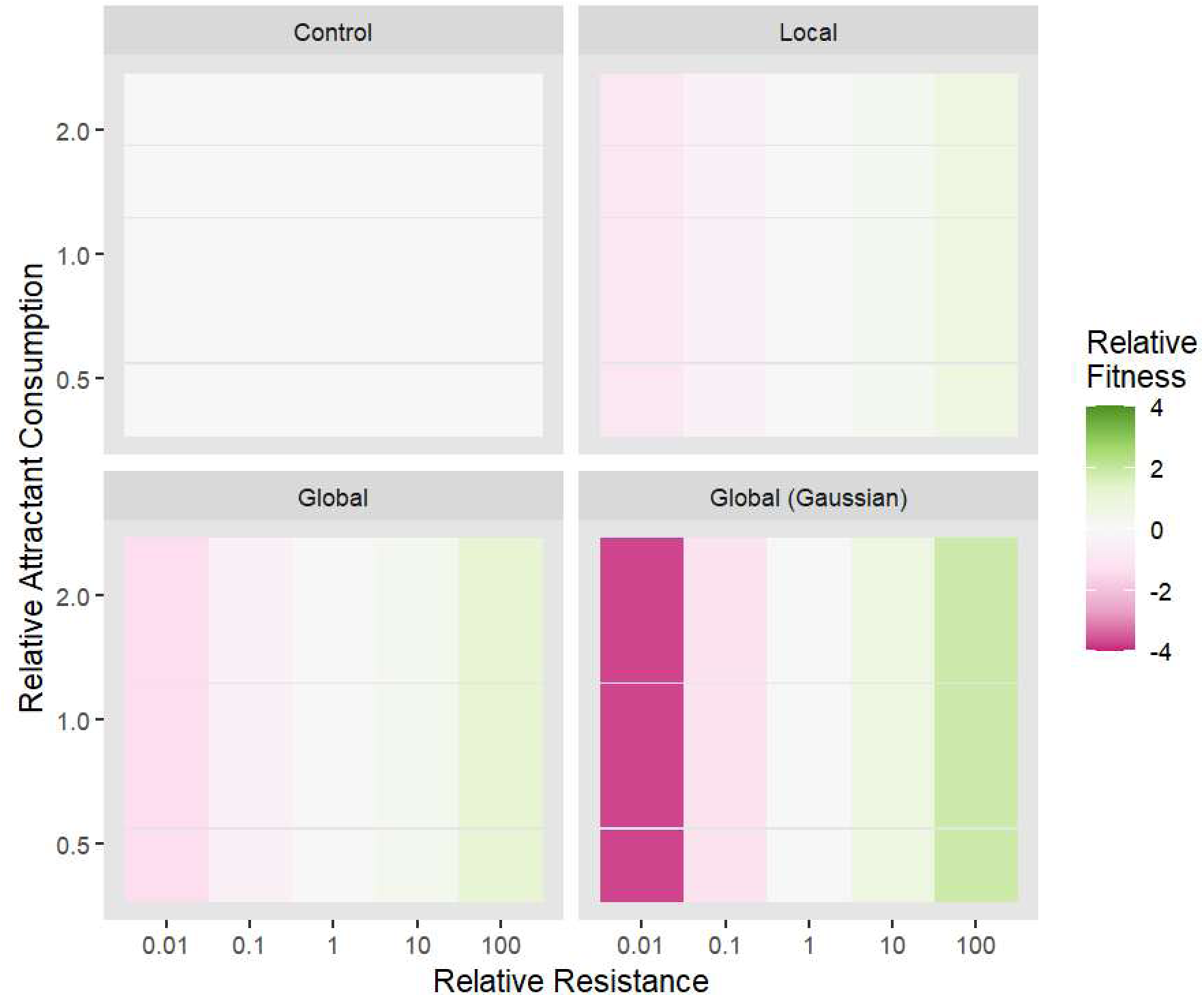
Parasite presence selects for host resistance with no effect on selection for host attractant consumption. *In silico* invasion experiments were carried out with a bacterial mutant that varied in its traits relative to the resident bacterial population, both competing in the presence of one of four different initial parasite spatial distributions. Results are colored by invader fitness = log_10_(final invader frequency/final resident frequency). Resistance is defined as the reciprocal of infection rate (i).

**Fig S12.**
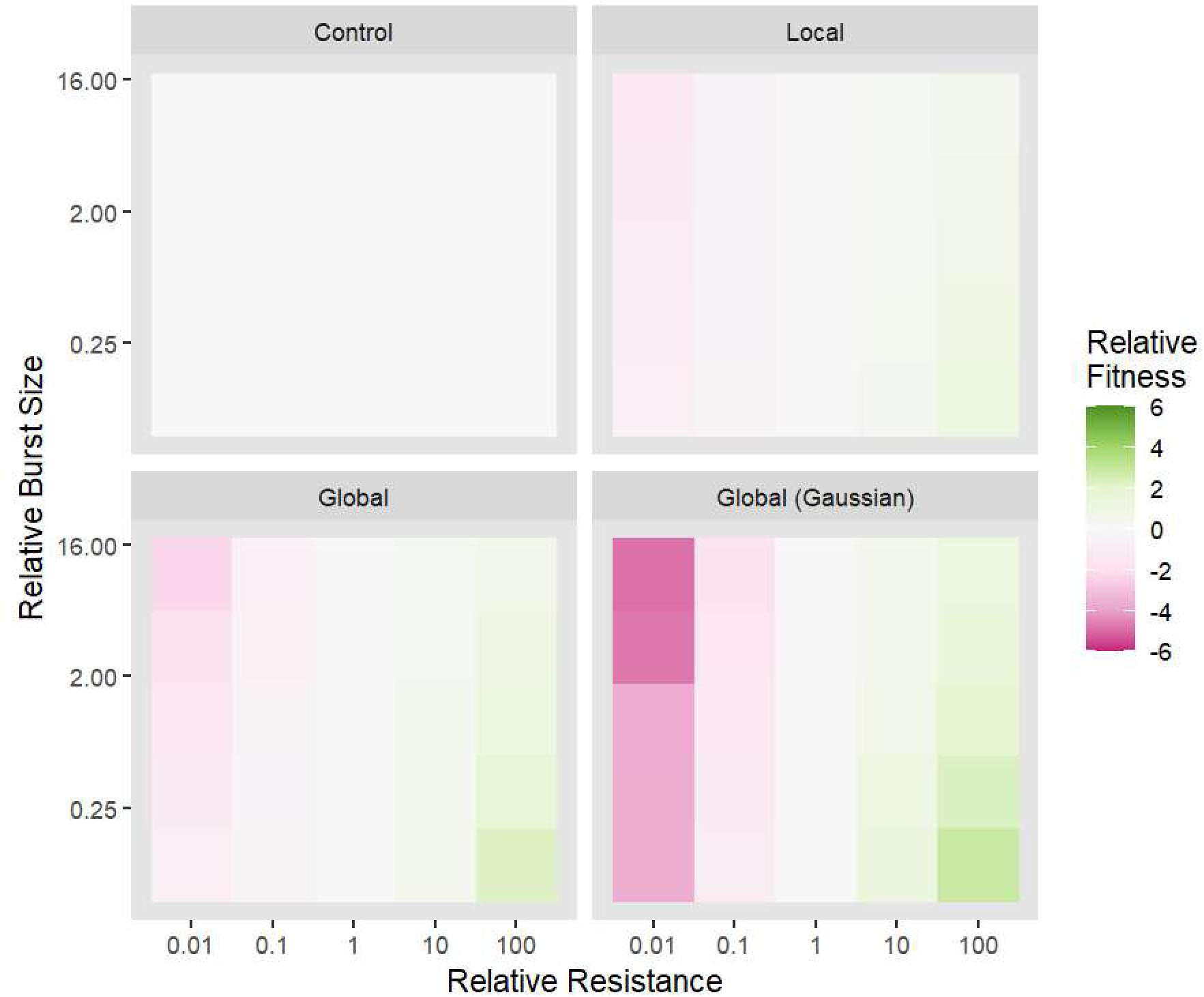
Parasite presence selects for host resistance but only weakly on burst size. *In silico* invasion experiments were carried out with a bacterial mutant that varied in its traits relative to the resident bacterial population, both competing in the presence of one of four different initial parasite spatial distributions. Results are colored by invader fitness = log_10_(final invader frequency/final resident frequency). Resistance is defined as the reciprocal of infection rate (i).

**Fig S13.**
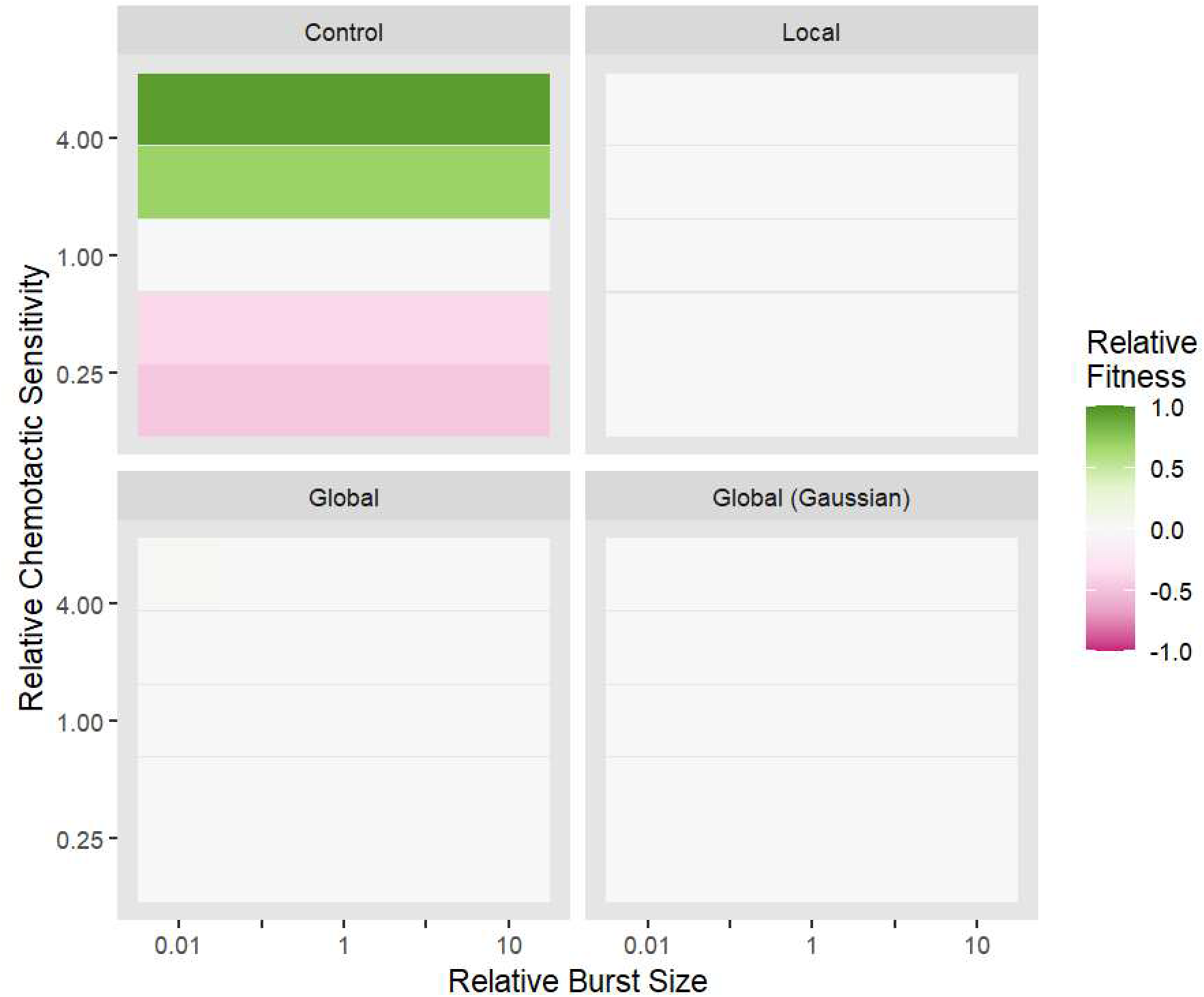
Chemotactic sensitivity is beneficial in the absence of parasites, but burst size is never beneficial. *In silico* invasion experiments were carried out with a bacterial mutant that varied in its traits relative to the resident bacterial population, both competing in the presence of one of four different initial parasite spatial distributions. Results are colored by invader fitness = log_10_(final invader frequency/final resident frequency).

**Fig S14.**
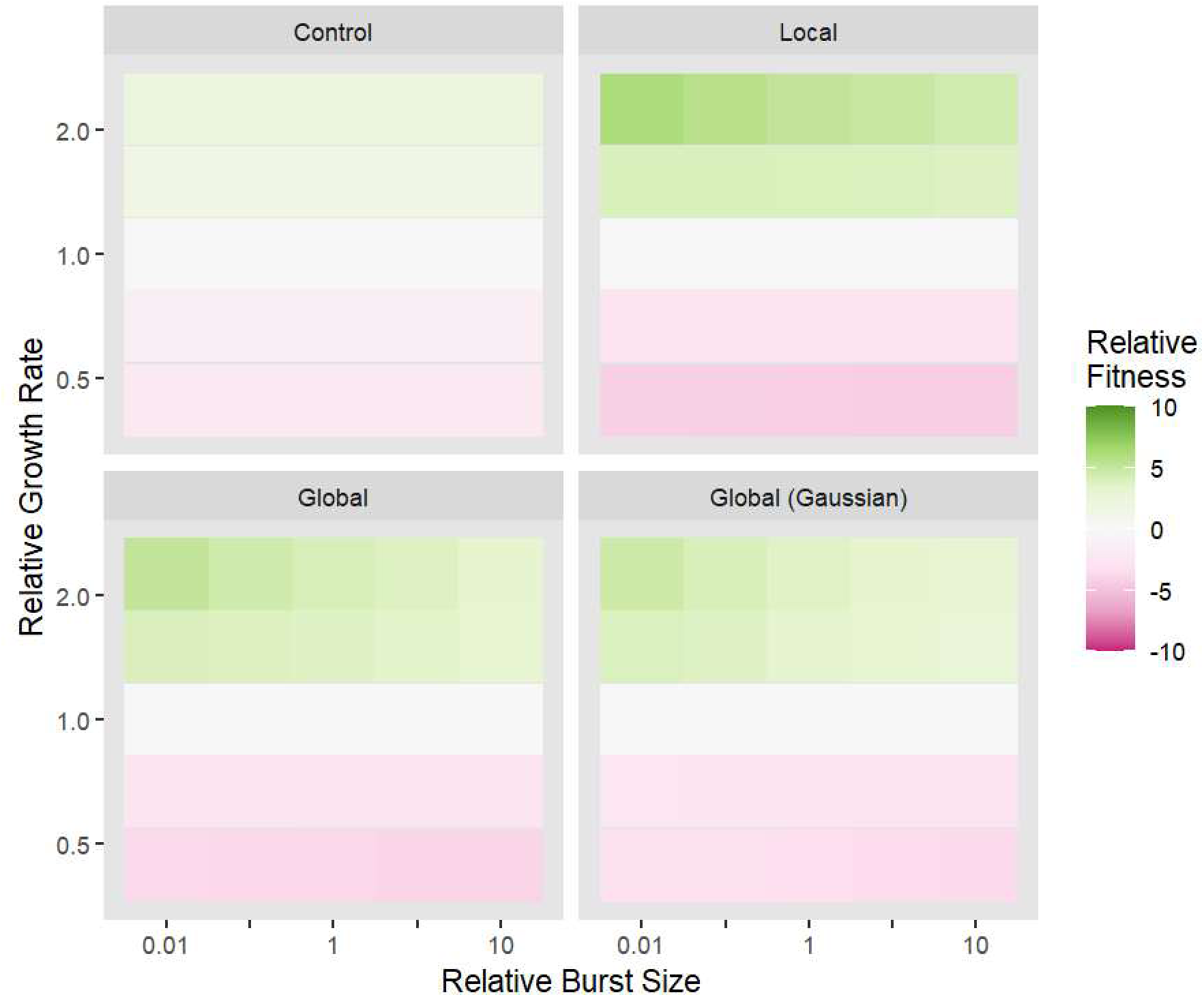
Growth rate is always beneficial, but burst size is weakly beneficial. *In silico* invasion experiments were carried out with a bacterial mutant that varied in its traits relative to the resident bacterial population, both competing in the presence of one of four different initial parasite spatial distributions. Results are colored by invader fitness = log_10_(final invader frequency/final resident frequency).

**Fig S15.**
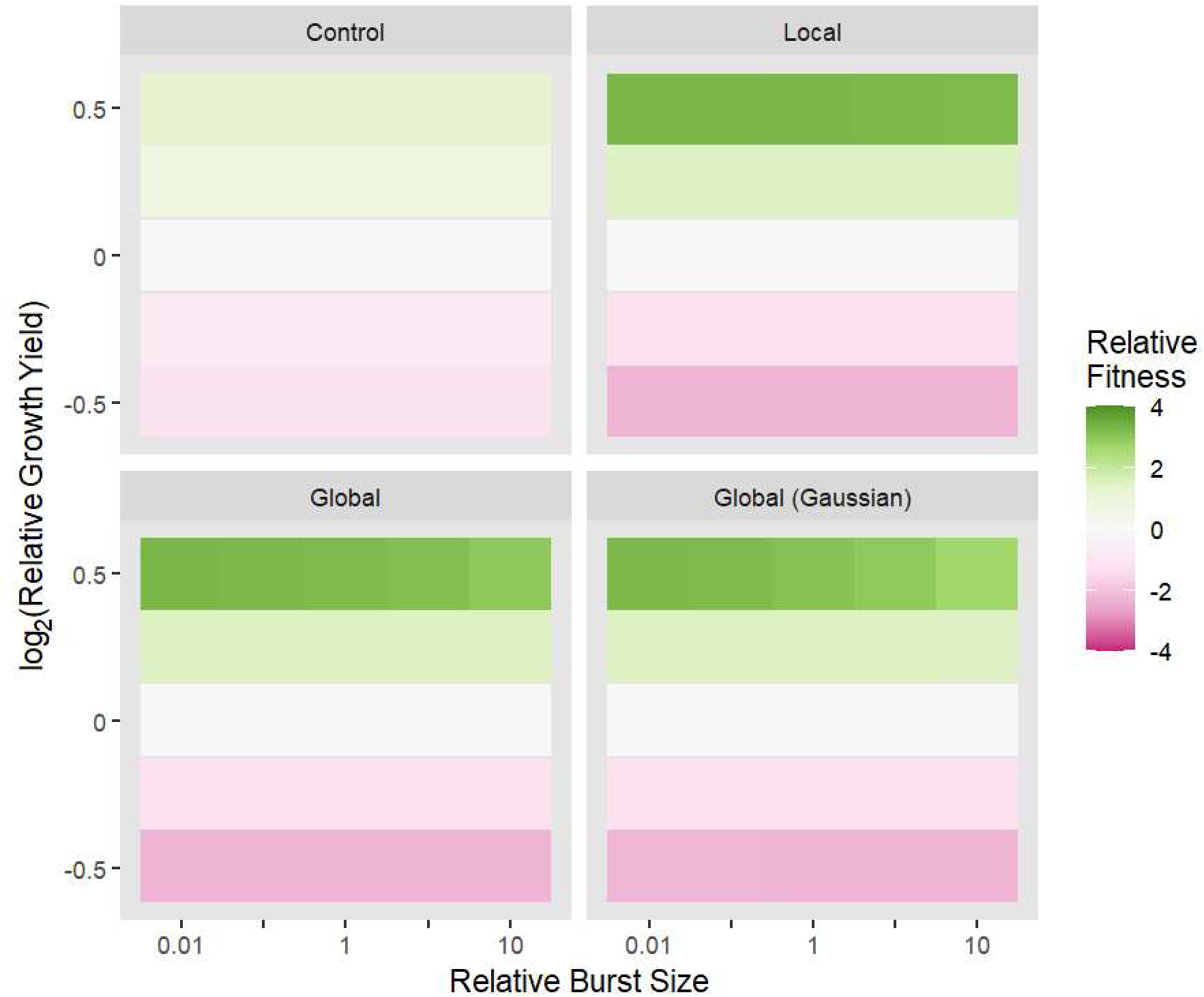
Yield is always beneficial while burst size is barely beneficial. *In silico* invasion experiments were carried out with a bacterial mutant that varied in its traits relative to the resident bacterial population, both competing in the presence of one of four different initial parasite spatial distributions. Results are colored by invader fitness = log_10_(final invader frequency/final resident frequency).

**Fig S16.**
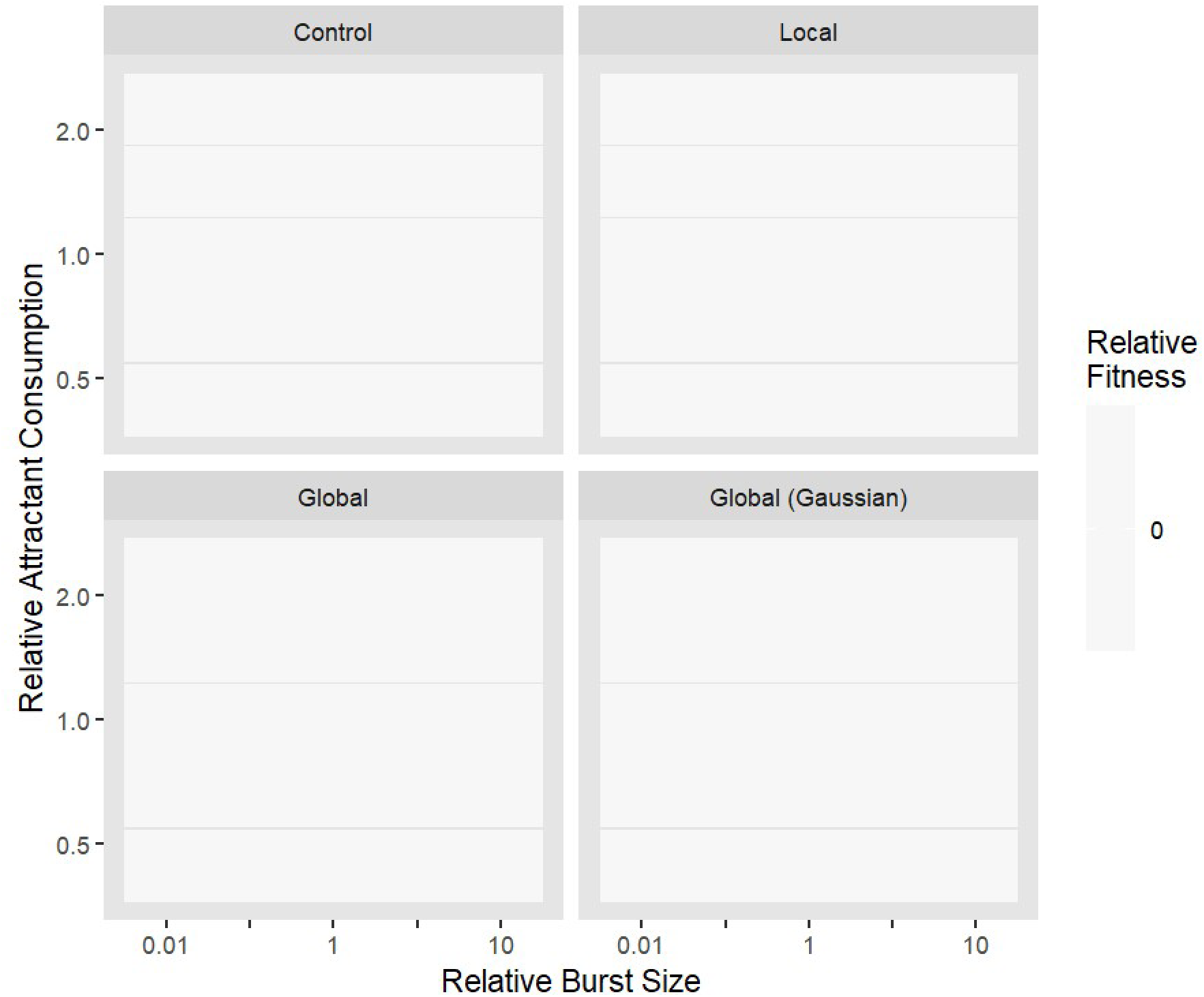
Neither attractant consumption nor burst size are beneficial. *In silico* invasion experiments were carried out with a bacterial mutant that varied in its traits relative to the resident bacterial population, both competing in the presence of one of four different initial parasite spatial distributions. Results are colored by invader fitness = log_10_(final invader frequency/final resident frequency).

**Table S2.** Statistical tests for differences in maximum per-capita growth rate. Mixed effects modeling of maximum per-capita growth rate was carried out using lmer. As a null model, we included random effects for the population and for the batch date. We then compared this null model to models with fixed effects with a Kenward-Roger approximate F-test. We included a fixed effect for treatment [ancestor, control, local, or global], a fixed effect for resistance [none, partial (EOP above detection limit but below 0.1), or complete (EOP below detection limit)], or both. No tests were significant, indicating that neither treatment nor resistance were significantly associated with evolved maximum per-capita growth rate in either condition or media.

| Experiment | Media | Fixed Effects | Approximate F-Statistic | p-value |
| --- | --- | --- | --- | --- |
| Less favorable for phage growth | Original | Treatment | 1.57 | 0.26 |
| Less favorable for phage growth | Rich | Treatment | 2.64 | 0.09 |
| Less favorable for phage growth | Original | Resistance | 2.49 | 0.11 |
| Less favorable for phage growth | Rich | Resistance | 2.56 | 0.11 |
| Less favorable for phage growth | Original | Treatment + Resistance | 2.96 | 0.07 |
| Less favorable for phage growth | Rich | Treatment + Resistance | 1.30 | 0.33 |
| More favorable for phage growth | Original | Treatment | 2.03 | 0.16 |
| More favorable for phage growth | Rich | Treatment | 1.25 | 0.34 |
| More favorable for phage growth | Original | Resistance | 0.80 | 0.46 |
| More favorable for phage growth | Rich | Resistance | 0.35 | 0.71 |
| More favorable for phage growth | Original | Treatment + Resistance | 1.24 | 0.34 |
| More favorable for phage growth | Rich | Treatment + Resistance | 0.72 | 0.62 |

**Table S3.** Parameters and values used in simulations. The sources for each parameter value and range of parameter values are listed. The fitting or derivation of some values is listed in more detail below.

| Parameter | Description | Resident Value | Invader Values | Source |
| --- | --- | --- | --- | --- |
| $D_N$ | Diffusion coefficient for bacteria | $50 \mu\text{m}^2/\text{s}$ | N/A | (Cremer et al. 2019) |
| $D_P$ | Diffusion coefficient for phage | $0 \mu\text{m}^2/\text{s}$ | N/A | Below |
| $D_R$ | Diffusion coefficient for resource | $800 \mu\text{m}^2/\text{s}$ | N/A | (Cremer et al. 2019) |
| $D_A$ | Diffusion coefficient for attractant | $800 \mu\text{m}^2/\text{s}$ | N/A | (Cremer et al. 2019) |
| $\chi$ | Chemotactic sensitivity coefficient | $315 \mu\text{m}^2/\text{s}$ | $57 - 1770 \mu\text{m}^2/\text{s}$ | (Cremer et al. 2019) |
| $c_R$ | Uptake rate of resource | $2.25 \times 10^{-14} \text{ mmol/cfu/s}$ | $1 - 5 \times 10^{-14} \text{ mmol/cfu/s}$ | Fitted, below |
| $c_A$ | Uptake rate of attractant | $4.17 \times 10^{-16} \text{ mmol/cfu/s}$ | $1.8 - 9.4 \times 10^{-16} \text{ mmol/cfu/s}$ | (Cremer et al. 2019) |
| $Y$ | Cell yield | $1.229 \times 10^{10} \text{ cfu/mmol}$ | $0.873 - 1.733 \times 10^{10} \text{ cfu/mmol}$ | Fitted, below |
| $K_R$ | Michaelis-Menten constant for resource | $5 \times 10^{-5} \text{ mmol/mL}$ | N/A | (Cremer et al. 2019) |
| $K_A$ | Michaelis-Menten constant for attractant | $1 \times 10^{-6} \text{ mmol/mL}$ | N/A | (Cremer et al. 2019) |
| $i$ | Infection rate | $1 \times 10^{-12} \text{ infec/cell/mL/pfu/mL/s}$ | $0.01 - 100 \times 10^{-12} \text{ infec/cell/mL/pfu/mL/s}$ | (Paepe and Taddei 2006; Wang 2006; Heineman and Bull 2007; Shao and Wang 2008; Gallet et al. 2009; Turner et al. 2012; Lindberg et al. 2014) |
| $b$ | Number of parasites produced per infection | $30 \text{ pfu/infec}$ | $3 - 300 \text{ pfu/infec}$ | (Josslin 1970; Reader and Siminovitch 1971; Paepe and Taddei 2006; Wang 2006; Heineman and Bull 2007; Gallet et al. 2009; Shao and Wang 2009; Turner et al. 2012; Lindberg et al. 2014) |

